# FGF-21 Conducts a Liver-Brain-Kidney Axis to Promote Renal Cell Carcinoma

**DOI:** 10.1101/2023.04.12.536558

**Authors:** Zongyu Li, Xinyi Zhang, Wanling Zhu, Cuiling Zhang, Katherine Sadak, Alexandra A. Halberstam, Jason R. Brown, Curtis J. Perry, Azia Bunn, David A. Braun, Adebowale Adeniran, Sangwon Lee, Andrew Wang, Rachel J. Perry

## Abstract

Metabolic homeostasis is one of the most exquisitely tuned systems in mammalian physiology. Metabolic homeostasis requires multiple redundant systems to cooperate to maintain blood glucose concentrations in a narrow range, despite a multitude of physiological and pathophysiological pressures. Cancer is one of the canonical pathophysiological settings in which metabolism plays a key role. In this study, we utilized REnal Gluconeogenesis Analytical Leads (REGAL), a liquid chromatography-mass spectrometry/mass spectrometry-based stable isotope tracer method that we developed to show that in conditions of metabolic stress, the fasting hepatokine fibroblast growth factor-21 (FGF-21)^1, 2^ coordinates a liver-brain-kidney axis to promote renal gluconeogenesis. FGF-21 promotes renal gluconeogenesis by enhancing β2 adrenergic receptor (Adrb2)-driven, adipose triglyceride lipase (ATGL)-mediated intrarenal lipolysis. Further, we show that this liver-brain-kidney axis promotes gluconeogenesis in the renal parenchyma in mice and humans with renal cell carcinoma (RCC). This increased gluconeogenesis is, in turn, associated with accelerated RCC progression. We identify Adrb2 blockade as a new class of therapy for RCC in mice, with confirmatory data in human patients. In summary, these data reveal a new metabolic function of FGF-21 in driving renal gluconeogenesis, and demonstrate that inhibition of renal gluconeogenesis by FGF-21 antagonism deserves attention as a new therapeutic approach to RCC.

## Main Text

Renal cell carcinoma (RCC), a prevalent cancer responsible for approximately 15,000 deaths each year in the U.S., is positively associated with obesity^3, 4^, highlighting the possibility that metabolic changes may drive RCC progression. Indeed, isotope tracing reveals a shift toward glycolytic and away from oxidative metabolism in human RCC tumors^5^. These data strongly hint at the potential for metabolism-targeting interventions to revolutionize treatment of RCC. However, none of the current therapeutic approaches to RCC – surgery, checkpoint inhibitors, tyrosine kinase inhibitors, and mTOR inhibitors – primarily alter metabolism, with the exception of a glutaminase inhibitor, which recently failed to improve outcomes in patients with metastatic RCC^6, 7^. Taken together, these data suggest that renal cell carcinoma is in great need of new metabolic approaches specifically targeting glucose metabolism to improve outcomes in RCC.

The ability of glucose to fuel tumor growth is not a new concept. In the 1920s, Otto Warburg popularized the idea that glucose promotes tumor growth by allowing the cell to meet its energetic demands via glycolysis. This shift away from oxidative metabolism shunts carbons to generate the biomass required to meet the demands of rapidly dividing tumor cells, including nucleotides, phospholipids, and amino acids. The tumor metabolism world’s canonical focus on glycolytic versus oxidative metabolism, however, ignores a central question: what is the source of the glucose that is avidly taken up by tumor cells? In a gluconeogenic organ such as the kidney, it would be logical that local increases in glucose production play a critical role in fueling tumor growth. In considering factors that could link renal glucose production to RCC, fibroblast growth factor-21 (FGF-21) emerges as a potential candidate. Knott and colleagues found that patients with RCC exhibited three-fold higher serum FGF-21 concentrations than healthy controls, and that high serum FGF-21 was an independent predictor of worse survival^8^. However, whether FGF-21 is simply a biomarker of poor prognosis in RCC, or plays a direct role in RCC pathology is unknown. Further, to our knowledge, FGF-21 has not been targeted for the treatment of RCC or any other tumor.

FGF-21 is a liver-derived fasting hormone^9, 10^. The metabolic role of FGF-21 has been best appreciated as a chronic activator of energy expenditure and therefore, of insulin sensitivity^11–19^. However, the idea that a fasting hormone’s sole metabolic effect could be catabolic, increasing energy dissipation during a period when the organism should be conserving energy, is unconvincing. This apparent paradox inspired us to examine the potential anabolic effects of FGF-21 during metabolic stress.

The role of the kidney in defending against metabolic stress remains an open question, in large part due to methodologic limitations. Renal glucose production has been approximated in humans by measuring arterio-venous differences in blood glucose concentrations and in plasma glucose enrichment during a tracer infusion. However, even using the same method and similar, healthy, recently fed subjects, renal glucose production has been reported in a strikingly wide range, between 0^20^ and 30%^21^ of total gluconeogenesis. Additionally, the methods typically used to measure renal glucose production in humans require cannulation of the renal artery and vein. This method is invasive and unlikely to be possible in mice, as would be necessary to permit knockout studies to facilitate mechanistic exploration of the metabolic role of the kidney in (patho)physiology. This discrepancy highlights the need for developing and validating alternative approaches to measure renal glucose production, and applying these improved approaches in the setting of RCC, in order to generate new metabolic targets for this devastating disease.

To address these unmet needs, we adapted our Positional Isotopomer NMR Tracer Analysis (PINTA) method^22^ to distinguish renal from hepatic glucose production^23^ using REGAL. Longstanding controversies exist as to whether arterial infusion and venous sampling, or venous infusion and arterial sampling, is optimal for *in vivo* tracer studies^24, 25^. However, in this *in vivo* setting, endogenous glucose production measured with arterial infusion and venous sampling did not differ from that measured with venous infusion and arterial sampling (Extended Data Fig. 1A). We thus infused tracer systemically through a jugular venous catheter advanced into the right atrium in subsequent studies. We also performed several validation studies to ensure that the REGAL method detected expected differences in renal glucose production in the setting of physiologically predictable alterations. First we infused glycerol, which promotes gluconeogenesis proportional to the dose infused in a largely unregulated manner^26^. As anticipated, we found that glycerol increased both hepatic and renal glucose production, without altering the fractional contribution of the kidney to whole-body glucose metabolism (Extended Data Fig. 1B-E). Next, we treated mice with a small molecule inhibitor of glycogen phosphorylase, which is expected to inhibit hepatic but not renal glucose production because the kidneys do not release significant glucose into circulation from glycogenolysis^27^. As expected, REGAL analysis demonstrated a reduction in plasma glucose and insulin concentrations, and in hepatic but not renal glucose production (Extended Data Fig. 1F-I).

**Figure 1.**
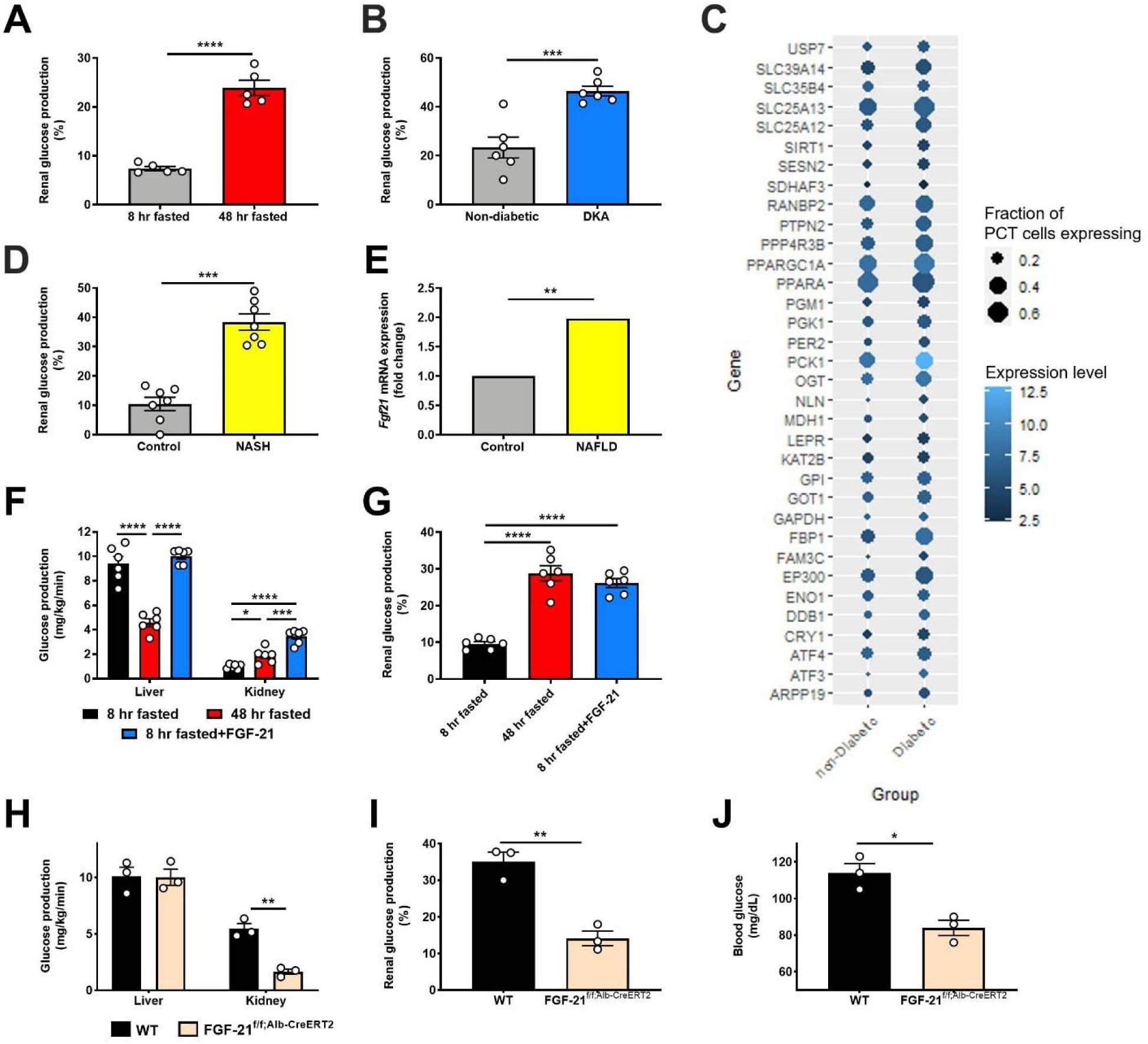
FGF-21 promotes renal gluconeogenesis under conditions of metabolic stress. (A)-(B) Renal glucose production increases in fasting (n=5 per group) and diabetic ketoacidosis (n= 6 per group) in mice. (C) Renal gluconeogenic gene expression increases in humans with diabetic nephropathy (n=3 per group). (D) Renal glucose production increases in a mouse model of NASH. (E) Liver *Fgf21* expression in humans with NAFLD (n=72, vs. n=6 healthy controls). (F)-(G) Recombinant FGF-21 infusion increases renal glucose production in rats (n=6 per group). (H)-(I) Hepatic and renal glucose production in 24 hour fasted liver-specific FGF-21 KO mice (n=3 per genotype). (J) Blood glucose concentrations (n=3 per genotype). In all panels, \**P*<0.05, \*\**P*<0.01, \*\*\**P*<0.001, \*\*\*\**P*<0.0001 by the 2-tailed unpaired Student’s t-test (panels A-B, D, H-J) or by ANOVA with Tukey’s multiple comparisons test (panels F-G). *P*-values for gene expression (panel E) were adjusted for multiple comparisons.

Having validated the REGAL method, we applied it to directly measure renal gluconeogenesis under conditions of metabolic stress of varying etiologies, both hypo- and hyper-caloric. Renal glucose production increased during a prolonged fast and in diabetic ketoacidosis (DKA) in mice (Fig. 1A-B, Extended Data Fig. 1J-Q). When we utilized a previously published dataset (Gene Expression Omnibus [GEO], Accession Number GSE131882) in humans with diabetic nephropathy, we found that expression of genes in the gene ontology pathway related to gluconeogenesis also increased in proximal tubule cells (Fig. 1C). Similarly, in mice with nonalcoholic steatohepatitis (NASH), we observed an increase in renal glucose production (Fig. 1D, Extended Data Fig. 1R-X).

Observing that, as expected, renal glucose production increased under disparate conditions of metabolic stress, we next asked what drove this adaptive response. We observed an increase in plasma FGF-21 in fasted mice, mice in DKA, and mice with NASH, as well as an increase in liver *Fgf21* gene expression in humans with non-alcoholic fatty liver disease (NAFLD) (GEO, Accession Number GSE130970, Fig. 1E, Extended Data Fig. 1Y-AA). We hypothesized that this protein may be responsible for the increased renal glucose production. To test this hypothesis, we performed a 2 hr infusion of recombinant FGF-21 to increase plasma FGF-21 concentrations in 8 hr fasted mice to match those measured in 48 hr fasted animals (Extended Data Fig. 1BB). Although FGF-21 had no effect on hepatic glucose production, it tripled both rates of renal glucose production and the fractional contribution of the kidney to whole-body glucose turnover (Fig. 1F-G, Extended Data Fig. 1CC-DD).

To confirm the source and role of FGF-21 without potential confounders from exogenous FGF-21 infusion, we generated liver-specific FGF-21 knockout (FGF-21^f/f;Alb-Cre^) mice. In contrast to their WT littermates, in which fasting increased plasma FGF-21 concentrations, FGF-21^f/f;Alb-Cre^ mice showed undetectable FGF-21 after a 24 hr fast (Extended Data Fig. 1EE). After a fast, FGF-21^f/f;Alb-Cre^ mice exhibited minimal renal glucose production and were consequently hypoglycemic with undetectable plasma insulin concentrations (Fig. 1H-J, Extended Data Fig. 1EE-FF). This demonstrates that FGF-21 is a liver-derived signal that upregulates renal glucose production during a fast.

These results were initially surprising in light of previous data demonstrating that chronic infusion of exogenous FGF-21 and constitutive FGF-21 overexpression reduces body fat content and improves glucose metabolism in animals with diet-induced obesity^11–19, 28, 29^. Consistent with previous studies, we observed an increase in energy expenditure, reductions in ectopic lipid content, and improved insulin sensitivity in mice with diet-induced obesity, as reflected by reduced *ad lib* fed plasma insulin concentrations in high fat fed mice (Extended Data Fig. 2A-L). However, when diet-induced obese mice underwent a prolonged (48 hr) fast, chronic FGF-21 infusion improved the ability of mice to maintain blood glucose concentrations.

**Figure 2.**
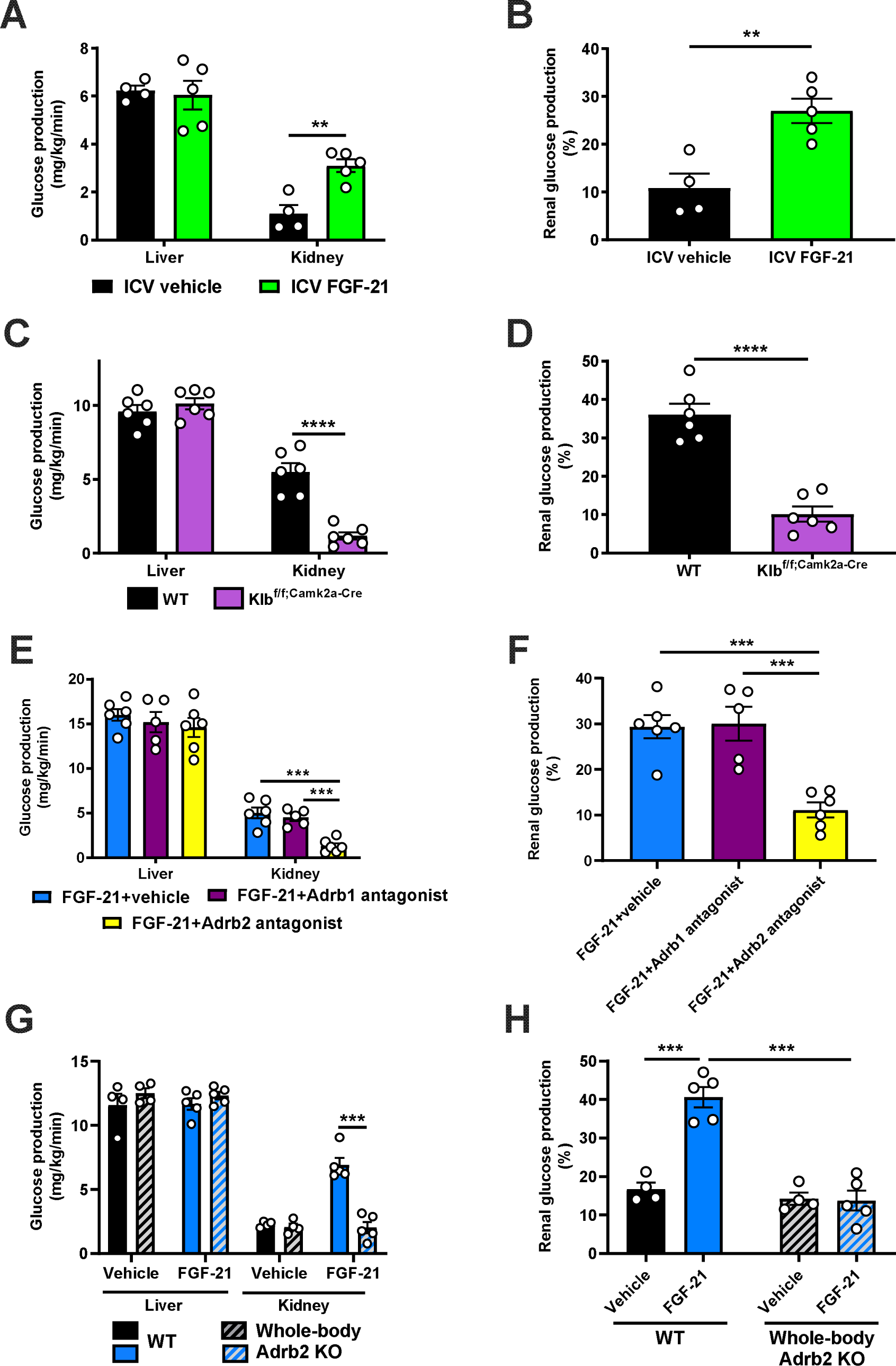
FGF-21 promotes renal gluconeogenesis via Adrb2-dependent neural hardwiring. (A)-(B) Endogenous glucose production from liver and kidney, and the fractional contribution of the kidney to total glucose production in rats administered an ICV infusion of FGF-21 (n=4 vehicle and 5 FGF-21). (C)-(D) Endogenous glucose production, and the renal contribution to whole-body glucose production in Klb^f/f;Camk2a-Cre^ mice (n=6 per genotype). (E)-(F) Endogenous glucose production, and the renal contribution to whole-body glucose production in 6 hr fasted mice infused with FGF-21 and pretreated with an Adrb1 or Adrb2 antagonist, or vehicle (n=6 vehicle, 5 Adrb1 antagonist, and 6 Adrb2 antagonist). (G)-(H) Endogenous glucose production, and the renal contribution to whole-body glucose production in Adrb2 knockout mice infused with FGF-21 (n=4 vehicle-treated in both genotypes, and 5 FGF-21-treated in both genotypes). In all panels, \*\**P*<0.01, \*\*\**P*<0.001, \*\*\*\**P*<0.0001 by the 2-tailed unpaired Student’s t-test.

Next, we sought to determine the mechanism by which FGF-21 enhances renal glucose production. As FGF-21 is known to regulate energy expenditure centrally^12, 30–32^, we hypothesized that its promotion of renal glucose production may also be centrally mediated. Consistent with this hypothesis, intracerebroventricular (ICV) FGF-21 injection increased renal glucose production despite unaltered circulating plasma FGF-21 concentrations (Fig. 2A-B, Extended Data Fig. 2M-O).

We then examined the impact of a prolonged fast on systemic glucose metabolism in mice lacking the FGF-21 coreceptor, β-Klotho (KLB), in the brain (Klb^f/f;Camk2a–Cre^ mice)^32, 33^, where both the FGF-21 receptor FGFR1c and Klb are highly expressed, unlike in liver or kidney^34, 35^. After a 48 hr fast, despite robust FGF-21 production, mice with brain-specific Klb deletion failed to induce renal, but not hepatic, glucose production. Consequently, fasted Klb^f/f;Camk2a–Cre^ mice were hypoglycemic, despite close to undetectable plasma insulin concentrations (Fig. 2C-D, Extended Data Fig. 2P-R). These data confirm that FGF-21 enhances renal glucose production and defends against hypoglycemia during a fast via a central action.

Next, we aimed to delineate the mechanism by which FGF-21 promotes renal glucose production. As β-adrenergic activity was previously shown to be critical for the effect of FGF-21 to cause adipose browning^30^, we first employed 6-hydroxydopamine (6-OHDA) to induce chemical sympathectomy. 6-OHDA indeed prevented FGF-21’s ability to stimulate renal glucose production, while also lowering hepatic glucose production (Extended Data Fig. 2S-W). We then examined the impact of a nonselective β-adrenergic antagonist, propranolol, on FGF-21-stimulated renal glucose production. We found that propranolol prevented the effect of FGF-21 to enhance renal gluconeogenesis (Extended Data Fig. 2X-BB).

Having demonstrated that FGF-21 promotes renal glucose production through central β-adrenergic activity, we then applied pharmacologic agonists and knockout models to further delineate which β-adrenergic receptor (Adrb) was responsible for this effect. As Adrb3 is not expressed in the kidney^35^, we applied Adrb1 and Adrb2 antagonists (betaxolol and butoxamine, respectively). We generated Adrb1 and Adrb2 knockout mice to validate these agonists. Adrb1 antagonism did not alter epinephrine-stimulated renal glucose production in Adrb1 knockout mice, and Adrb2 antagonism did not alter epinephrine-stimulated renal glucose production in Adrb2 knockout mice (Extended Data Fig. 2CC). Having demonstrated the specificity of the antagonists, we applied them in mice infused with FGF-21. Whereas the Adrb1 antagonist had no impact on FGF-21-stimulated renal gluconeogenesis, the Adrb2 antagonist fully abrogated the ability of FGF-21 to promote renal glucose production (Fig. 2E-F, Extended Data Fig. 2DD-FF). To conclusively demonstrate the role for Adrb2 in FGF-21-driven renal gluconeogenesis, we infused FGF-21 in whole-body Adrb2 knockout (KO) mice and their WT littermates. FGF-21 failed to stimulate renal glucose output in Adrb2 KO mice, while it increased both renal glucose production, plasma glucose and insulin concentrations in WT animals (Fig. 2G-H, Extended Data Fig. 2GG-II).

Finally, to determine to what extent circulating catecholamines mediate the impact of FGF-21 to stimulate renal glucose production, we studied fed and 24 hour fasted adrenalectomized (ADX) mice. To dissociate their lack of glucocorticoid production from any phenotype observed in ADX mice, we infused ADX mice with corticosterone by subcutaneous pellet. This ensured that fasting plasma corticosterone concentrations were matched between sham-operated and ADX animals. Under these conditions, we observed no differences in plasma glucose and insulin concentrations, or renal glucose production, between sham-operated and ADX mice after a 24 hour fast (Extended Data Fig. 2JJ-OO). Taken together, these data demonstrate that Adrb2-mediated neural hardwiring underlies FGF-21’s effect to stimulate renal glucose production in mice.

Finally, we sought to determine how Adrb2 signaling promotes renal gluconeogenesis. We previously demonstrated that glucagon stimulates hepatic gluconeogenesis by enhancing intrahepatic lipolysis^36^. We hypothesized that a similar mechanism may underlie FGF-21’s effect on the kidney. Consistent with this hypothesis, we observed an increase in kidney long-chain acyl- and acetyl-CoA concentrations without accompanying increases in whole-body lipolysis (palmitate turnover) (Fig. 3A-B, Extended Data Fig. 3A), which, together, reflects increased intrarenal lipolysis. Acetyl-CoA is an allosteric activator of pyruvate carboxylase (PC)^37, 38^ but its role in renal gluconeogenesis has remained unknown. Consistent with a role for acetyl-CoA-mediated regulation of renal glucose production, we observed increased kidney PC activity in mice infused with FGF-21 (Fig. 3C). However, each of these effects of FGF-21 on readouts of intrarenal lipolysis and PC activity were prevented by Adrb antagonism with propranolol. We confirmed increased intrarenal lipolysis – as reflected by increases in kidney long-chain acyl-CoA and acetyl-CoA concentrations – in each of the metabolic stress models tested: fasting, DKA, NASH, and ICV FGF-21 infusion. We found that brain-specific Klb knockout, chemical sympathectomy, Adrb2 antagonism, and Adrb2 knockout fully abrogated the effect of FGF-21 to stimulate intrarenal lipolysis (Fig. 3D-E, Extended Data Fig. 3B-W).

**Figure 3.**
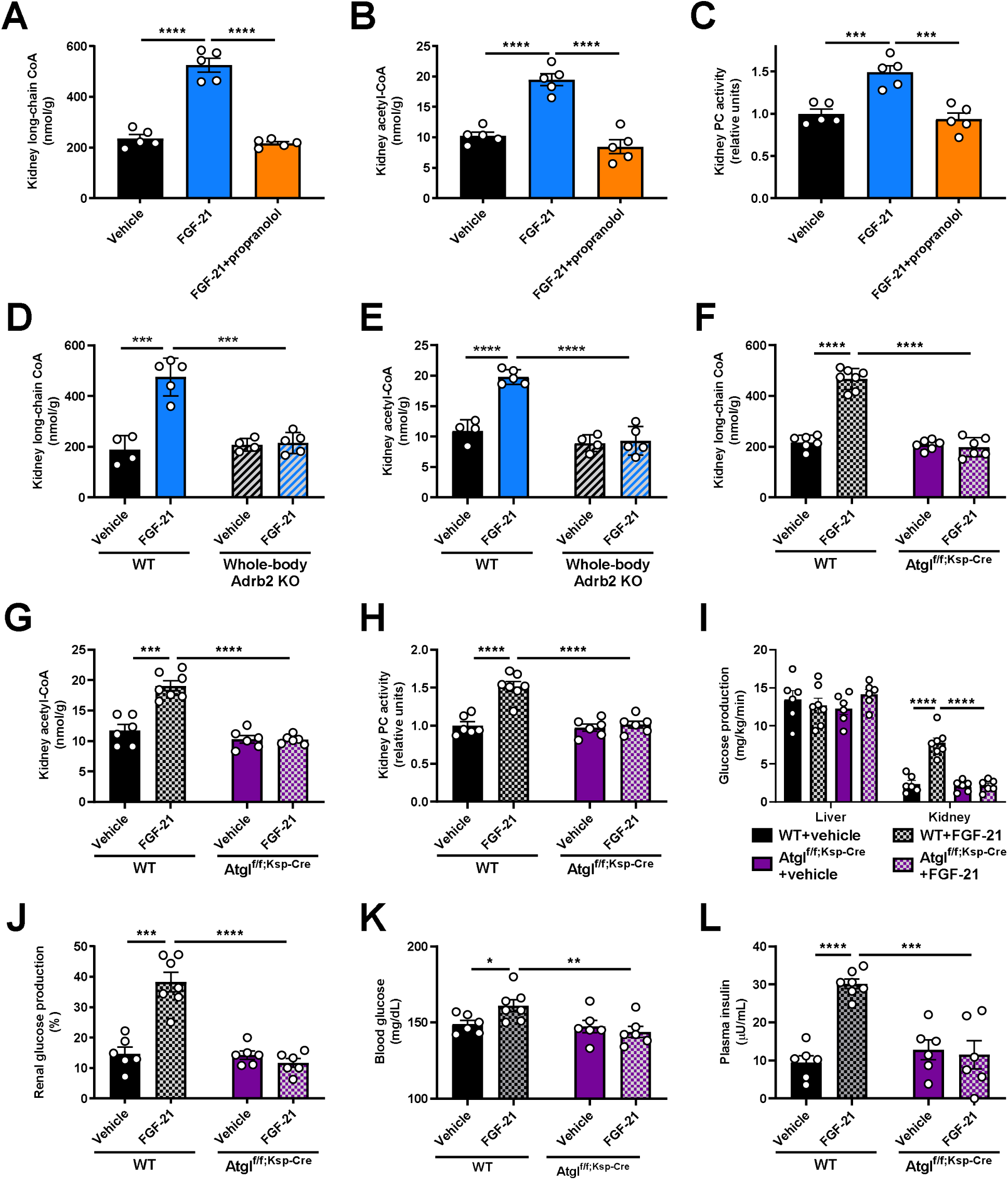
FGF-21-induced increases in renal lipolysis promote increased renal gluconeogenesis in metabolic stress. (A)-(B) Kidney long-chain acyl- and acetyl-CoA concentrations in mice infused with FGF-21±the nonspecific Adrb antagonist propranolol. In panels (A)-(C), groups were compared by ANOVA with Tukey’s multiple comparisons test, and n=5 per group. (C) *Ex vivo* pyruvate carboxylase (PC) activity. (D)-(E) Kidney long-chain acyl- and acetyl-CoA concentrations in whole-body Adrb2 knockout mice (n=4 vehicle-treated and 5 FGF-21-treated per genotype). In panels (D)-(L), groups were compared by the 2-tailed unpaired Student’s t-test. (F)-(G) Kidney long-chain acyl- and acetyl-CoA concentrations in kidney-specific ATGL knockout mice (Atgl^f/f;Ksp-Cre^) (n=6 per group with the exception of WT+FGF-21-treated mice [n=7 per group]). (H) Kidney pyruvate carboxylase activity (in panels (H)-(L), n=6 per group with the exception of WT+FGF-21-treated mice [n=7 per group]). (I)-(J) Endogenous glucose production, and the renal contribution to whole-body glucose production. (K)-(L) Blood glucose and plasma insulin concentrations. In all panels, \**P*<0.05, \*\**P*<0.01, \*\*\**P*<0.001, \*\*\*\**P*<0.0001.

To conclusively confirm the effect of FGF-21 to stimulate renal gluconeogenesis via intrarenal lipolysis, we generated collecting duct-specific ATGL knockout (Atgl^f/f;Ksp-Cre^) mice. We chose this segment of the nephron because it contains the highest levels of expression of *Atgl* mRNA^35^. In contrast to WT mice, in which FGF-21 again enhanced intrarenal lipolysis, PC activity, and gluconeogenesis, Atgl^f/f;Ksp-Cre^ mice did not exhibit any increase in kidney lipolysis, PC activity, or gluconeogenesis in response to FGF-21 (Fig. 3F-L, Extended Data Fig. 3X). These data demonstrate that under metabolic stress, as a result of increased intrarenal lipolysis and acetyl-CoA content, FGF-21 enhances PC activity to promote renal gluconeogenesis.

Considering the clear ability of FGF-21 to enhance renal glucose production, we next hypothesized that FGF-21 could be a novel target for renal cell carcinoma. Indeed, we observed a progressive increase in plasma FGF-21 concentrations in three well-established murine models of kidney cancer: B6.129S4-Tsc2^tm1Djk/J^ mice, a murine model of tuberous sclerosis in which renal adenomas progressively develop between 6 and 16 months of age^39^; Six2^CreERT^^2^;Vhl^f/f^;Bap1^+/-^ mice with induced RCC^40^; and WT mice with RCC (Renca) cells injected into the renal medulla^41^ and metastasizing to lung (Extended Data Fig. 4A). In each model, plasma FGF-21 increased as tumors developed (Fig. 4A-C). The FGF-21 was not derived from the tumor: human RCC tumors contained negligible FGF-21 mRNA, and mouse kidneys with and without RCC did not contain measurable FGF-21 protein (Extended Data Fig. 4B-C). However, in mice injected with Renca cells that did not grow a tumor (likely due to immune suppression of tumor progression), FGF-21 induction was reduced by more than 90% (Extended Data Fig. 4D). Taken together, these data demonstrate that the presence of RCC induces FGF-21 production in mice.

**Figure 4.**
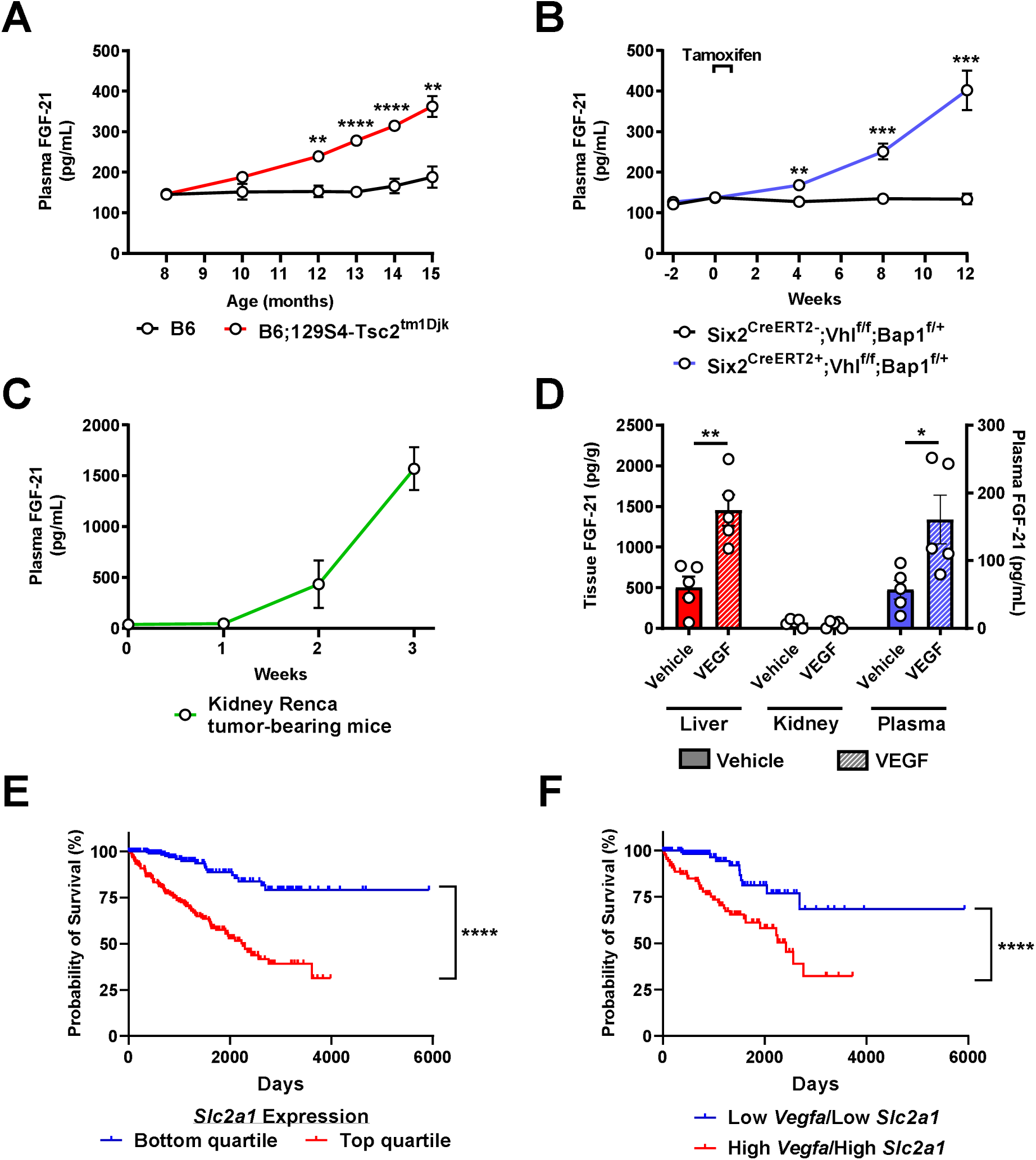
FGF-21 increases in a VEGF-dependent manner in murine models of kidney cancer. (A) Plasma FGF-21 concentrations in a mouse model of renal adenoma (n=5 per group). (B)-(C) Plasma FGF-21 concentrations in mouse models of renal cell carcinoma (n=5 per group, with the exception of week 3 in kidney Renca tumor-bearing mice, in which n=4 due to the death of one of the mice). (D) FGF-21 concentrations in healthy mice treated acutely with recombinant VEGF (n=5). (E) Survival probability in RCC patients with low and high (lowest and highest 25^th^ percentile) *Slc2a1* (GLUT1) expression^43^ (n=219 per group). (F) Survival probability in RCC patients with low and high (lowest and highest 25^th^ percentile) expression of both *Vegfa* (VEGF) and *Slc2a1* (GLUT1)^43^ (n=224 per group). In all panels, \**P*<0.05, \*\**P*<0.01, \*\*\**P*<0.001, \*\*\*\**P*<0.0001.

Next we aimed to determine the mechanism of induction of FGF-21 in response to kidney cancer in mice. We reasoned that the signal for FGF-21 production must be a protein produced by the RCC tumor. The canonical growth factor produced by RCC tumors is the pro-angiogenic vascular endothelial growth factor (VEGF), which is a poor prognostic factor in patients with RCC (Extended Data Fig. 4E). VEGF concentrations increased seven-fold in mice with kidney Renca tumors, whereas we observed a much smaller increase was observed in mice injected with Renca cells that did not grow visible tumors (Extended Data Fig. 4F-G). Injection of VEGF induced FGF-21 protein in liver and in plasma, but not in kidney (Fig. 4D), and also modestly induced adipose tissue lipolysis reflected by an increase in plasma non-esterified fatty acid (NEFA) concentrations (Extended Data Fig. 4H), as has been shown previously^42^.

The combination of the demonstrated ability of FGF-21 to promote renal gluconeogenesis and the prognostic effect of the glucose transporter GLUT1 in highlight the potential for VEGF-driven FGF-21 in promoting RCC progression via activating renal glucose production. Consistent with a critical role for glucose in promoting RCC progression, high expression of *Slc2a1*, which encodes the primary tumor glucose transporter, GLUT1, is strongly associated with worse survival in patients with RCC: data from the Human Protein Atlas show a ten-year survival rate of 79% in patients in the lowest quartile of RCC *Slc2a1* expression, as compared to just 39% in patients in the top quartile (Fig. 4E)^43^. Patients in the lowest quartile of both *Vegfa* and *Slc2a1* expression fared even worse, with a ten-year survival rate of just 32% (Fig. 4F).

These data suggest that FGF-21-driven renal gluconeogenesis is a targetable pathogenic factor downstream of VEGF in RCC progression. We treated Renca tumor-bearing mice with an Fc-fused FGF-21 C-terminal peptide to block the activity of endogenous FGF-21 by masking ligand-binding sites on KLB^44^. After a 48 hr fast, mice treated with the peptide were hypoglycemic, associated with markedly reduced renal glucose production as compared to mice treated with PBS vehicle (Fig. 5A-C, Extended Data Fig. 5A-B). Consistent with prior data demonstrating an effect of FGF-21 to reduce bile acid synthesis, inactivating FGF-21 with the Fc-fused FGF-21 c-terminal peptide also increased the total bile acid concentration in plasma (Extended Data Fig. 5C). While we cannot inject Renca cells (BALB/c background) into our complement of knockout mice on a C57bl/6J background, we performed *ex vivo* studies to ascertain the mechanism by which FGF-21 drives kidney glucose production in the setting of RCC. Consistent with the idea that tumors are net glucose consumers while kidney can be a net glucose producer as directed by FGF-21 under conditions of metabolic stress, when we examined tumors and surrounding kidney parenchyma in mice with kidney Renca tumors, we observed a marked increase in net glucose production in normal kidneys as compared to tumors.

**Figure 5.**
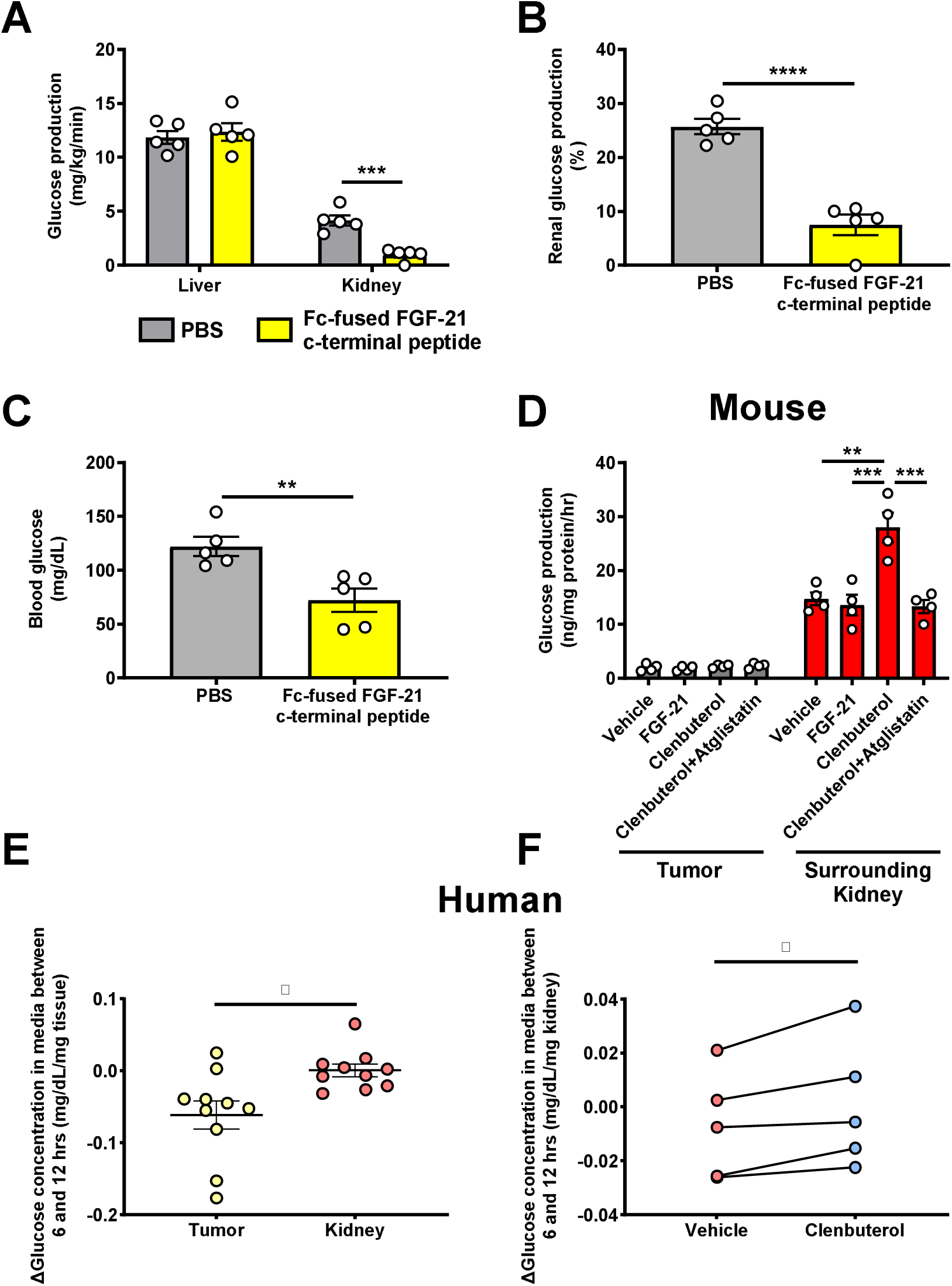
FGF-21-dependent Adrb2 activity promotes renal glucose production in mice with RCC. (A)-(C) Glucose production and blood glucose concentrations in Renca tumor-bearing mice treated with a Fc-fused FGF-21 c-terminal peptide (n=5 per group). (D) Glucose production in murine Renca and surrounding kidney samples (n=4 tumor and 4 surrounding kidney). (E) Glucose production in human RCC tumor and surrounding parenchyma (n=10). (F) Glucose production in human RCC parenchyma treated with clenbuterol or vehicle (n=5). In panels (E) and (F), the paired t-test was used because samples from the same patients were compared. In all panels, \**P*<0.05, \*\**P*<0.01, \*\*\**P*<0.001, \*\*\*\**P*<0.0001.

Next, we examined the impact of the Adrb2 agonist on renal glucose production *ex vivo.* Net glucose production was accelerated by Adrb2 activity and dependent upon intrarenal lipolysis in normal kidney parenchyma: the Adrb2 agonist clenbuterol accelerated glucose production, but atglistatin, an inhibitor of the key lipolytic enzyme adipose triglyceride lipase, abrogated the effect of clenbuterol to promote kidney parenchyma glucose production. In contrast, recombinant FGF-21 had no effect to increase kidney glucose production, consistent with our hypothesis that the central nervous system mediates FGF-21’s effect to accelerate renal gluconeogenesis through Adrb2 (Fig. 5D).

Next, we sought to determine the translatability of these results to human patients. Kidney parenchyma from patients undergoing nephrectomy for RCC showed higher net glucose production under basal conditions in both normal parenchyma and tumors. (Fig. 5E, Extended Data Table 1). Treatment with the Adrb2 agonist clenbuterol stimulated renal glucose production in non-tumor kidney parenchyma but not tumor, (Fig. 5F). Taken together, these data are consistent with a critical role for high net gluconeogenesis driven by *Adrb2* and *Atgl* activity in the surrounding renal parenchyma and low net gluconeogenesis in the avidly glucose-utilizing tumor. These data again suggest that inhibiting the FGF-21-Adrb2-renal gluconeogenesis axis holds promise for treatment of RCC.

Finally, we sought to use these new insights into the metabolic regulation of RCC via a VEGF-FGF-21-Adrb2-ATGL-gluconeogenesis axis to generate new therapeutic approaches. Considering that Adrb2 activity mediated the metabolic effects of FGF-21 in kidney in mouse and human, we hypothesized that treatment with a nonselective Adrb2 blocker, propranolol, would slow tumor growth in mice with Renca RCC. Indeed, we observed that chronic propranolol treatment reduced tumor size in kidney Renca tumor-bearing mice (Fig. 6A-B). Strikingly, observational data indicate that survival in human patients with RCC at two institutions treated with the nonselective Adrb blocker propranolol is lower than in RCC patients not treated with propranolol (Fig. 6C), although with the caveat that we were not able to control for many confounders including RCC subtype, clinicopathologic characteristics, tumor stage, or other medications.

**Figure 6.**
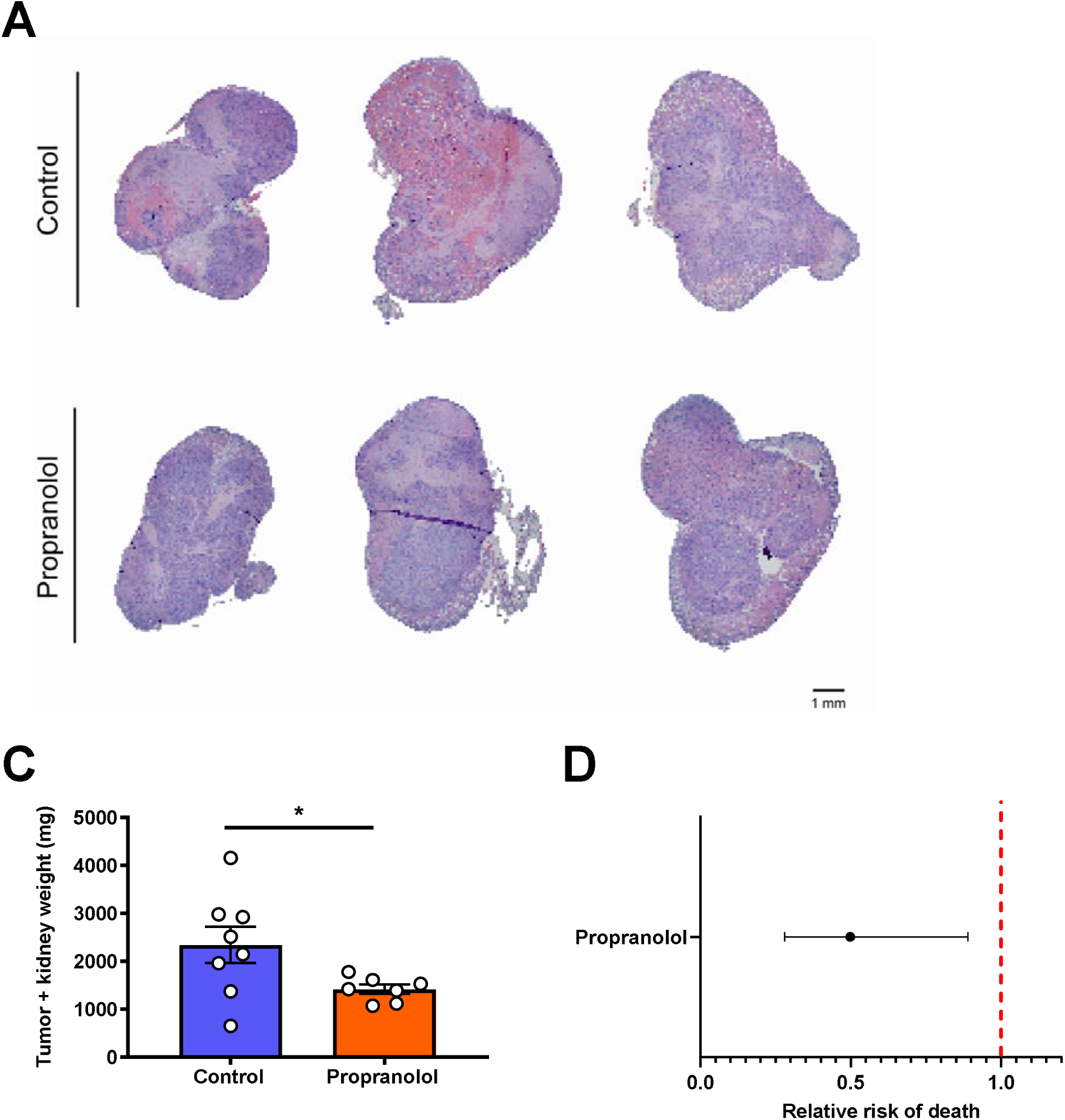
Increased renal glucose production promotes RCC tumor progression. (A) Representative images of kidney stained with hematoxylin & eosin from Renca tumor-bearing mice treated with propranolol. (B) Treatment with the Adrb antagonist propranolol slows Renca tumor progression in mice (n=8 controls and 7 propranolol-treated mice). \**P*<0.05. (C) RCC patients who have ever been prescribed propranolol have improved survival as compared to RCC patients who have never taken propranolol. The 95% confidence interval is shown.

Data from the Human Protein Atlas show that patients with high expression of the glucose transporter *Slc2a1* and low expression of the gluconeogenic enzyme *Pck1* (cytosolic phosphoenolpyruvate carboxykinase) in their RCC tumors have worse survival than those with low *Slc2a1* and high *Pck1* expression (28% vs. 70% survival at 3,000 days) (Extended Data Fig. 6A)^43^. Similarly, patients with low *Adrb2* and low *Atgl* expression in tumors had worse survival than those with high *Adrb2* and high *Atgl* expression (Extended Data Fig. 6B). Comparing tumors with low expression of *Adrb2, Atgl*, and *Pck1* to tumors with high expression of *Adrb2, Atgl*, and *Pck1* was underpowered (27 patients with low expression of all three proteins, and 14 patients with high), but strongly suggested a survival benefit in patients with low expression of all three key proteins in the pathway connecting FGF-21 to renal glucose production (Extended Data Fig. 6C). However, it is important to note that these are data in RCC tumors, not the surrounding kidney parenchyma: our data would imply that high gluconeogenic protein expression in kidney parenchyma, but not tumors (which are net glucose consumers) would predict poor outcomes. Taken together, the data in this study suggest that FGF-21 or Adrb2 inhibition may be attractive therapeutic strategies for patients with RCC, due to their expected effect to inhibit renal gluconeogenesis.

While the kidney has long been known to have the capacity for gluconeogenesis, its role in metabolic homeostasis has been underappreciated due to a paucity of methods capable of assessing renal glucose production that can be applied in rodents. A recent study reinvigorated interest in renal gluconeogenesis using stable isotope tracers^45^ but did not aim to elucidate the mechanism of regulation of renal glucose production under conditions of metabolic stress. Here we apply novel REGAL tracer methodology (Extended Data Fig. 7A-B), seven transgenic mouse models, six targeted pharmacologic approaches, and four surgical approaches – to demonstrate that FGF-21- and Adrb2-mediated intrarenal lipolysis increases kidney gluconeogenesis, and that this FGF-21-dependent kidney gluconeogenesis axis promotes RCC (Extended Data Fig. 7C).

FGF-21 has been appreciated for its ability to increase energy expenditure, reduce ectopic lipid content, and enhance systemic glucose metabolism. These data reveal an a formerly unappreciated role for FGF-21 in coordinating the gluconeogenic response to metabolic stress, and demonstrate that the metabolic role of FGF-21 is substantially larger than previously thought. Not only does FGF-21 play a role in counteracting the detrimental effects of overnutrition by enhancing energy expenditure, it also plays a surprising but crucial role in counteracting the detrimental effects of undernutrition by promoting renal glucose production under conditions of metabolic stress. These pleotropic effects of FGF-21 highlight the fact that the core function of this hormone is not well understood. Evolutionary pressures have generally selected for robust anabolic programs – to counteract hypoglycemia in fasting – substantially more than catabolic machinery. Tumors hijack these mechanisms. This study identifies approaches neutralizing FGF-21 or its downstream pro-gluconeogenic pathways as potential therapeutic targets against RCC. The development of new approaches to treat RCC is of particular interest because therapeutic options for invasive RCC are limited to checkpoint inhibitors and tyrosine kinase inhibitors. These agents improve progression-free survival by only several months and are associated with numerous adverse effects^46^. Therefore, the development of new therapeutic approaches is urgently needed. Here we identify a mechanism dependent on FGF-21 for the coordination of the systemic (tumor-liver-central nervous system-fat-kidney) response to the metabolic stress induced by renal cell carcinoma. Taken together, these data identify FGF-21-targeting therapies or Adrb2 blockade as a promising new therapeutic approach to renal cell carcinoma.

## Methods

### Data Availability

When possible (i.e. in all cases other than x-y and survival curves), all data are shown in the figures. All raw data generated *de novo* for this study are available in Supplementary File 1, without restrictions. For data generated by others, links/accession codes are provided.

### Code Availability

The code generated to analyze human *Fgf21* and gluconeogenic gene expression (Fig. 1C, E) is available at https://github.com/xz710/renal_gluconeogenesis.

### Animals

All procedures were approved by the Yale University Institutional Animal Care and Use Committee. All rodents used in these studies were male, Sprague-Dawley rats and mice on C57bl/6J or Balb/c background, between 8-14 weeks of age, with the exception of the Western diet fed mice as well as B6;129S4-Tsc2^tm1Djk^ and Six2^CreERT2+^;Vhl^f/f^;Bap1^f/+^ mice. C57bl/6J (stock number 000664), Balb/cJ (stock number 000651), and Adrb1/2 double knockout mice (stock number 003810) were purchased from Jackson Labs, and ADX mice (stock code ADREX) and sham-operated controls from Charles River. Upon arrival ADX mice underwent surgery to place subcutaneous pumps infusing corticosterone (2 mg/day) to match concentrations measured in fasted animals, thereby avoiding corticosterone as a physiologic confounder, and they were maintained on 0.9% sodium chloride drinking water. Adrb1 and Adrb2 mice were generated by backcrossing Adrb1/2 double knockout mice. Atgl^f/f;Ksp-Cre^ mice were generated by crossing Atgl^f/f^ and Ksp-Cre mice, both purchased from Jackson Labs. FGF-21^f/f;Alb-CreERT2^ mice were generated by crossing FGF-21^f/f^ (from Jackson Labs) and Alb^CreERT2^ mice (generously provided by Dr. Pierre Chambon to A.W.). In FGF-21^f/f;Alb-CreERT2^ mice, beginning two weeks prior to study, recombination was induced using tamoxifen (75 mg/kg/day in corn oil, IP, daily for five days). Klb^f/f^ and Camk2a-Cre mice were a kind gift from Dr. David Mangelsdorf and were used to generate Klb^f/f;Camk2a-Cre^ animals with brain-specific knockdown of the FGF-21 coreceptor^12, 32, 47^. Mice with NASH were purchased from Taconic on a 40% fat/20% fructose/2% cholesterol diet and were studied after 36 weeks on the diet following an overnight fast. Sprague-Dawley rats were purchased from Charles River, some with catheters in the third ventricle of the brain (stock code 3RDVENTCAN).

In all genetic models, genotyping was performed by quantitative PCR, using primers from IDT with the following sequences:

**Table 1.**
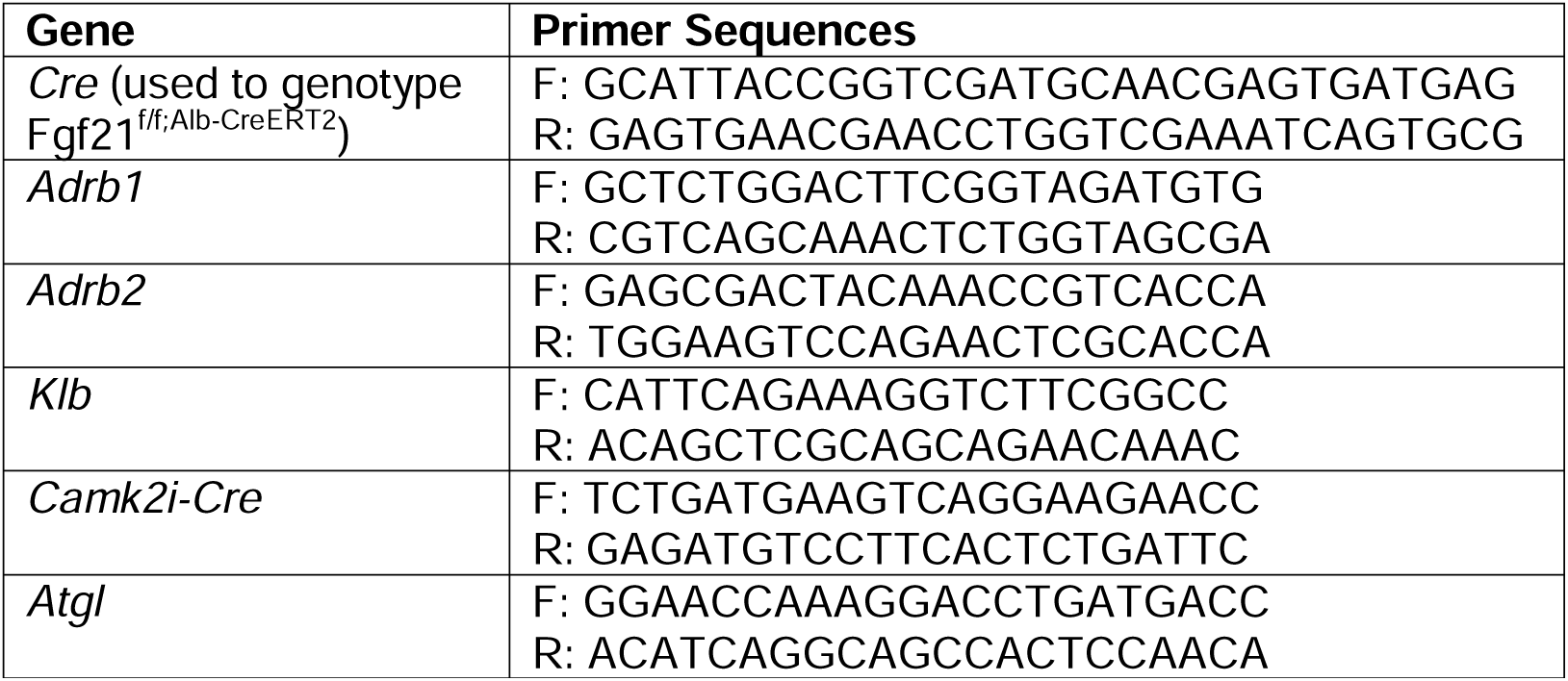
Primer sequences. used to genotype mice during the studies to delineate the mechanism of induction of renal gluconeogenesis in mouse models of metabolic stress.

Unless otherwise specified, mice were maintained on regular chow (Teklad #2018, 18% calories from fat, 24% from protein, 59% from carbohydrate) and drinking water *ad lib.* Mice were fasted for 8 or 48 hours prior to study, as indicated in the figures/figure legends. If not specified, a 8 hr fast was used.

Obesity was induced in a subset of mice by feeding C57bl/6J mice a high fat/high carbohydrate Western diet (*ad lib* access to a lard-based high fat diet containing 60% calories from fat/20% from protein/20% from carbohydrate, and 5% sucrose drinking water). Beginning on the first day of Western diet feeding, these mice were infused continuously with recombinant mouse FGF-21 (2.5 μg/day, similar to a previous report^48^), or PBS vehicle, by subcutaneous pump for four weeks. During week 1 of Western diet/FGF-21 infusion, before they diverged in body weight, mice underwent Comprehensive Lab Animal Monitoring System (Columbus Instruments) metabolic cage analysis to assess energetics, food and water consumption. Body fat content was measured by NMR (Bruker) after four weeks. Diabetic ketoacidosis was induced by injection of streptozotocin (200 mg/kg) in overnight fasted mice, 72 hr prior to a tracer study. Mice with severe hyperglycemia (≥250 mg/dL following an overnight fast) were included in the study.

### RCC Models

Six2^CreERT2^ (stock number 032488), Bap1^f/f^ (stock number 031565), and Vhl^f/f^ (stock number 004081) mice were purchased from Jackson Labs. Bap1^f/f^ and Vhl^f/f^ mice were backcrossed to generate Six2^CreERT2+^;Vhl^f/f^;Bap1^f/+^ mice and Cre-controls. Recombination was induced using tamoxifen (75 mg/kg/day in corn oil, IP, daily for five days, beginning at 8 weeks of age). Quantitative PCR was employed for genotyping, using the following primers from IDT:

**Table 2.**
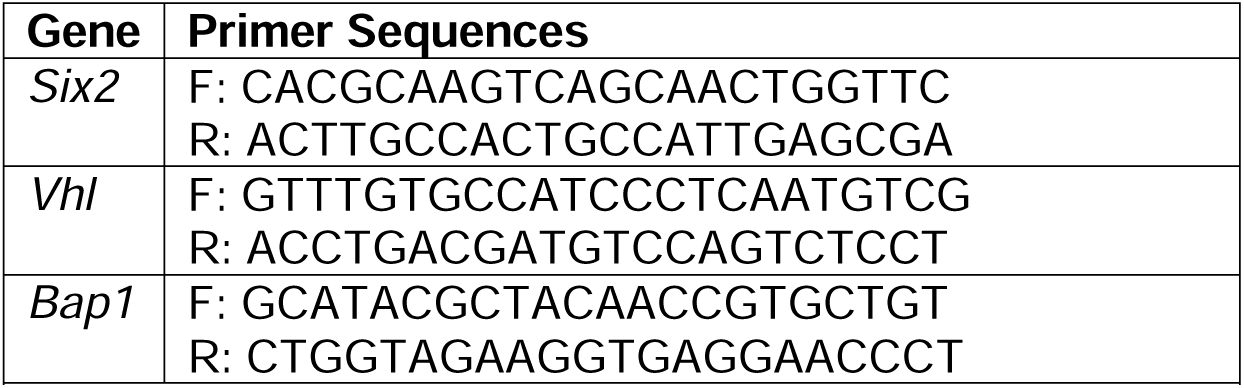
Primer sequences. used to genotype mice during the studies to delineate the mechanism by which FGF-21 promotes kidney cancer in murine models.

B6;129S4-Tsc2^tm1Djk^ mice (stock number 004686) were purchased from Jackson Labs. Tail vein plasma FGF-21 concentrations were measured monthly beginning at 8 months of age. Renca cells were purchased from ATCC and maintained in the manufacturer’s recommended media. Cells were not authenticated in our laboratory. After testing negative for mycoplasma contamination, 2x10^5^ Renca cells were injected into the kidney of wild-type Balb/c mice under isoflurane anesthesia as described by Murphy et al.^41^ Pharmacologic interventions as detailed below were applied in some mice, and plasma FGF-21 concentrations were measured in tail vein blood at the time points indicated in the figures. 21 days after tumor cell injection, mice were euthanized, and tumor (both in the kidney and the surrounding intraperitoneal space) weighed. Kidneys were fixed in 10% neutral buffered formalin, stained with hematoxylin & eosin by the Yale Comparative Medicine Research Histology Core, and examined by a blinded pathologist (author A.A.).

### Infusion Studies

Prior to flux studies, mice underwent surgery under isoflurane anesthesia to place catheters in the jugular vein advancing into the right atrium, and/or carotid artery. Rats underwent surgery to place catheters in the carotid artery, the jugular vein, advancing into the right atrium, and the descending aorta, advancing into both renal arteries.

After a week of recovery and following fasting for the duration specified, mice were placed in a plastic restrainer and tails gently taped in place, while rats remained unrestrained in their home cages. With exceptions described in the figure legends, tracers were infused into catheters in the jugular vein in mice and in the carotid artery in rats. Animals received a 120 min 3X primed-continuous infusion of [3-^13^C] lactate (Sigma; continuous infusion rate 20 μmol/kg/min for rats and 40 μmol/kg/min for mice) and [1,2,3,4,5,6,6-^2^H_7_] glucose (Cambridge Isotopes; continuous infusion rate 0.5 mg/kg/min for rats and 1.0 mg/kg/min for mice). In a subset of animals, [U-^13^C_16_] potassium palmitate (Cambridge Isotopes; continuous infusion rate 5 μmol/kg/min following a 3X prime) was infused concurrently to assess whole-body lipolysis. At the conclusion of the two hour infusion, blood was obtained from the tail vein (mice) or from the jugular vein (rats), centrifuged in heparin-lithium coated tubes, and plasma isolated. Animals were sacrificed using IV Euthasol (pentobarbital/phenytoin). Livers and kidneys were freeze-clamped in tongs pre-chilled in liquid nitrogen within 2 sec of excision (mice) or *in situ* (rats), and tissues were stored at -80°C pending further analysis.

### Pharmacologic Interventions

During a subset of tracer studies, FGF-21 was infused continuously into the jugular vein in mice, with catheters advancing into right atrium, the carotid artery in rats, (0.1 μg/hr or 1.0 μg/hr, respectively), or ICV over a ten min period at the beginning of a tracer infusion in rats (0.1 μg). An equivalent volume of PBS was administered to vehicle-treated animals. Chemical sympathectomy was achieved using 6-OHDA (10 mg/kg in PBS IP, 22 hr prior to the start of FGF-21 and tracer infusion). Antagonists of Adrb (propranolol, 5 mg/kg), Adrb1 (betaxolol, 5 mg/kg), and Adrb2 (butoxamine, 10 mg/kg), all from Sigma and dissolved in PBS, were administered IP, one hour prior to the start of FGF-21 and/or tracer infusion. Epinephrine (2 mg/kg total) was infused continuously, IV, throughout the tracer infusion in some mice. VEGF (1 mg/kg in PBS) was administered IP, and blood drawn from the tail vein 2 and 6 hr thereafter. For the tracer validation studies, glycogen phosphorylase was inhibited by IV (jugular vein) injection of 1-(3-(3-(2-Chloro-4,5-difluorobenzoyl)ureido)-4-methoxyphenyl)-3-methylurea (Sigma, 5 mg/kg) one hour prior to the start of a tracer infusion. In a separate group of mice, glycerol (50 μmol/kg/min) was infused continuously for 120 min concurrently with ^13^C lactate and ^2^H glucose, as described above (“Infusion Studies”).

The impact of manipulating Adrb2 activity on RCC progression was tested in mice with Renca tumor cells in the kidney. Mice were randomized to receive propranolol (0.5 mg/mL, approximate daily dose 30 mg/kg) in drinking water, or control water not containing propranolol, beginning 5 days before tumor cell injection so as to allow mice to acclimate to the drinking water. A construct for Fc-fused FGF-21 C-terminal peptide (Fc-FGF-21_CT_) was generated by ligating a DNA sequence corresponding to a signal peptide from murine heavy chain, Fc region of human IgG1, a (GGGGS)_2_ linker, followed by C-terminal region of human FGF-21 (amino acids 166-209) into a vector pCEP4 (Thermo Fisher Scientific). The construct was then transiently transfected to Expi293F cells (Thermo Fisher Scientific) following the protocol from the manufacture. The media containing Fc-FGF-21_CT_ was harvested 4 days after the transfection. Fc-FGF-21_CT_ was purified using protein A sepharose 4B (Thermo Fisher Scientific) and dialyzed against PBS. Purified Fc-FGF-21_CT_ was subject to an endotoxin removal proces (Pierce High Capacity Endotoxin Removal Spin Columns, Thermo Fisher Scientific) before flash-frozen and stored at -80°C until use. Fc-FGF-21_CT_ was administered to Renca tumor-bearing mice (20 μg/mouse) 48 hr prior to a terminal study.

### REGAL Tracer Analysis

This method has been previously published^23^ but was validated and optimized and is therefore described in detail here. The tracer study workflow is shown in Extended Data Fig. 7A. Plasma ^13^C_7_ glucose enrichment was determined using gas chromatography/mass spectrometry (GC/MS)^49^ in the chemical ionization (CI) mode and used to calculate whole-body glucose turnover (Table 3, Equation 1) and whole-body gluconeogenesis (Equation 2).

**Table 3.** Flux ratios and absolute rates. measured in mice infused with [3-13C] lactate. APE indicates the atom percent enrichment, and GNG denotes gluconeogenesis.

| Equation | Interpretation |
| --- | --- |
| 1. $Turnover = \left( \frac{Tracer\ APE}{Plasma\ APE} - 1 \right) * Infusion\ rate$ | Whole-body endogenous glucose production (EGP) or palmitate turnover |
| 2. $Liver\ glycogenolysis = \frac{[Liver\ glycogen]_{t-2} - [Glycogen]_t}{t}$ | Liver glycogenolysis rate |
| 3. $Total\ gluconeogenesis = EGP - liver\ glycogenolysis$ | Whole-body gluconeogenesis (V<sub>GNG</sub>) |
| 4. $\frac{V_{PEPCK}}{V_{GNG}} \sim \frac{V_{PC}}{V_{GNG}} = \frac{[^{13}C_2]glucose}{XFE^2}$ | Fraction of gluconeogenesis derived from pyruvate, measured in plasma (i.e. whole-body), liver, or kidney |
| 5. $XFE = \frac{1}{1 + \frac{[^{13}C_1]_{glucose}}{2 \cdot [^{13}C_2]_{glucose}}}$ | Fractional triose enrichment |
| 6. $Corrected [^{13}C_2]_{glucose} = Measured [^{13}C_2]_{glucose} - 2 * [C4C5C6 - ^{13}C_2]_{glucose}$ | Doubly-labeled glucose arising from the condensation of two singly labeled trioses, correcting for doubly labeled glucose arising from one doubly labeled triose condensing with an unlabeled triose |
| 7. $\frac{GNG_K}{GNG} = 1 - \frac{GNG \text{ from pyruvate}_T - GNG \text{ from pyruvate}_K}{GNG \text{ from pyruvate}_L - GNG \text{ from pyruvate}_K}$ | Fractional contribution of the kidney to total gluconeogenesis |
| 8. $GNG_K = \frac{GNG_K}{GNG} * V_{GNG}$ | Absolute rate of gluconeogenesis from the kidney |

Plasma palmitate enrichment was measured by GC/MS (CI mode) and used to calculate turnover using the whole-body turnover equation^49^. Glycogen content was determined by the modified phenol-sulfuric acid method^50–52^. The rate of net hepatic glycogenolysis was assumed to be constant between n-2 and n hours of fasting^49^. By subtracting the rate of net hepatic glycogenolysis from the total rate of glucose turnover, we calculated the rate of whole-body gluconeogenesis (Table 3, Equations 2-3).

REGAL uses Mass Isotopomer Distribution Analysis (MIDA)^53^ to measure fractional glucose production from phosphoenolpyruvate (PEP) in plasma (reflecting whole-body gluconeogenesis from PEP), liver, and kidney. We calculated the fraction of glucose production from PEP in each of these tissues using MIDA (Table 3, Equations 4-5) with GC/MS measurement of ^13^C_1_ and ^13^C_2_ glucose enrichment in the CI mode. In these calculations, we correct for any ^13^C_2_ glucose synthesized from ^13^C_2_ trioses – as opposed to the condensation of two ^13^C_1_ trioses – by GC/MS measurement of the enrichment in the glucose C4C5C6 fragment, according to Equation 6 in Table 3. By comparing the whole-body rate of gluconeogenesis from pyruvate (GNG from pyruvate_T_) to that measured in liver (GNG from pyruvate_L_) and kidney (GNG from pyruvate_K_), we were able to measure the fractional contribution of the kidney to whole-body gluconeogenesis (Table 3, Equation 7). Absolute rates of glucose production from the kidney were determined by multiplying the fractional contribution of the kidney by the total endogenous glucose production rate (Table 3, Equation 8).

### Biochemical and Histological Analysis

Blood glucose concentrations were measured using a handheld glucometer (Auvon). Plasma FGF-21, insulin, corticosterone, and VEGF concentrations were measured by ELISA (R&D Systems, Mercodia, Abcam, and Sigma, respectively), and total plasma bile acids using a colorimetric assay (Abcam). Kidney PC activity was measured enzymatically^54^, and plasma NEFA using the Wako NEFA-HR(2) kit. Bicarbonate and transaminase (ALT, AST) concentrations were measured by COBAS, and tissue triglyceride concentrations measured enzymatically using the method of Bligh and Dyer^55^. Acetyl-^56^ and long-chain acyl-CoA concentrations^36^ were measured by LC-MS/MS while β-OHB by GC/MS^57^ as previously described. Sections of the right medial lobe of the liver were obtained from mice with NASH, fixed in 10% neutral buffered formalin, and stained with hematoxylin & eosin. Samples were examined and images captured by a blinded investigator using an Olympus BX51 multi-headed brightfield microscope (Yale Liver Center Morphology Core). Whole kidneys were obtained from mice with kidney cancer, fixed in 10% neutral buffered formalin, and stained with hematoxylin & eosin. Sections from the center of the kidney cortex were stained with hematoxylin & eosin, examined and images captured by a blinded investigator using an Olympus BX51 microscope.

### RNA Sequencing Analysis

Data from human patients with NAFLD^58^ (Gene Expression Omnibus, Accession Number GSE130970) and humans with diabetic nephropathy^59^ (Gene Expression Omnibus, Accession Number GSE131882) were compared to healthy controls using gene ontology enrichment analysis. Expression of *Vegfa* (available from https://www.proteinatlas.org/ENSG00000112715-VEGFA/pathology/renal+cancer), *Pck1* (available from https://www.proteinatlas.org/ENSG00000124253-PCK1/pathology/renal+cancer), *Slc2a1* (available from https://www.proteinatlas.org/ENSG00000117394-SLC2A1/pathology/renal+cancer), *Adrb2* (available from https://www.proteinatlas.org/ENSG00000169252-ADRB2/pathology/renal+cancer), *Atgl* (available from https://www.proteinatlas.org/ENSG00000177666-PNPLA2/pathology/renal+cancer), and *Fgf21* (available from https://www.proteinatlas.org/ENSG00000105550-FGF21/pathology/renal+cancer) was recorded and correlated to survival, using datasets in the Human Protein Atlas^43^. Considering the presence of 887 samples in the Human Protein Atlas database with mRNA expression and survival data, the top and bottom quartiles consisted of the 219 samples with the highest and 219 samples with the lowest expression of the proteins of interest, with the exception of tumor *Adrb2* expression. “Ties” (i.e. samples with expression equal to that of samples in the top or bottom quartile of expression of genes of interest) were all included in the data shown from the top or bottom quartiles.

### Survival Analysis in Patients with RCC

The EPIC SlicerDicer tool was employed to ascertain the fraction of patients in the Yale-New Haven Hospital System with RCC (ICD-10 C64) prescribed propranolol who were alive or deceased from when data accrual began in 1982, to when data were collected on 3/19/2023. Additionally, the EMERSE tool^60^ was utilized using keywords "renal cell carcinoma" and "propranolol" to review clinic notes and to ascertain the fraction of patients in the University Hospitals Cleveland system with RCC prescribed propranolol who were alive or deceased from when data accrual began in 2015 to data collection in 2023 under University Hospitals IRB approved protocol 20220322.

### Quantification and Statistical Analysis

Animals were randomized to treatment groups using Excel’s random number generator. Group sizes were predetermined based on a power calculation. GraphPad Prism 9 was used for all statistical analysis. Groups were compared by one-way ANOVA with Tukey’s multiple comparisons test (when comparing 3 or more groups), or by the 2-tailed paired or unpaired t-test (when comparing 2 groups), as denoted in the figure legends. Survival curves were compared using the log-rank (Mantel-Cox) test. Data are presented as the mean±S.E.M. No data were excluded from analysis. All data points shown are from biological replicates, not technical replicates. For all the flux studies and the measurements of β-OHB and transaminase concentrations, two technical replicates were analyzed per sample. Investigators were not blinded as to group allocation during the *in vivo* studies. However, all biochemical, histological, and flux analysis, as well as assessment of whether mice had reached humane endpoints, was performed by investigators who were blinded as to group allocations. No samples were excluded from analysis; however, in some cases, analyses were not performed due to either a failure in the tracer infusion study (as indicated by a mouse’s lack of response to IV euthanasia), lack of sufficient sample remaining to complete an analysis, or the death/euthanasia for humane endpoints of a mouse prior to the planned endpoint. A power calculation revealed that a sample size of 4 per group was expected to be sufficient based on an expected 50% difference in the key parameters of interest with a standard deviation of 25% (80% power, a=0.05); however, if statistical significance were achieved in the primary endpoint (renal glucose production) with n=3, then 3 samples were analyzed in some cases.

## Acknowledgments

We thank Dr. Pierre Chambon for kindly providing Alb^CreERT2^ mice, and Dr. David Mangelsdorf for generously supplying Klb^f/f^ and Camk2a-cre mice. This study was funded by grants from the U.S. Public Health Service (K99/R00 CA215315 [R.J.P.], R37 CA258261-01A1 [R.J.P.], K08 AI128745 [A.W.]), R01 AR080104 [A.W.], and from the Pew Charitable Trusts [A.W.]). The Yale Liver Center, through whose microscopy core we generated histology images from mice with NASH and from mice with renal cell carcinoma, is supported by P30 DK034989. We are grateful to RCC patients undergoing nephrectomy at Yale-New Haven Hospital for allowing us to study glucose production in their tumor and surrounding parenchyma.

## Author Contributions

The study was designed by R.J.P. and Z.L. Unique mouse models were generated and kindly provided by A.W. and C.Z., and human kidney samples by D.A.B. and K.S. Fc-fused FGF-21 c-terminal peptide was generously provided by S.L. Experiments were performed and data analyzed by Z.L., X.Z., W.Z., J.R.B., C.J.P., A.A., and R.J.P. The manuscript was drafted by Z.L. and R.J.P., and extensively edited by A.A.H., A.W., A.B., and X.Z. All authors reviewed and approved the final version before submission.

## Competing Interests Declaration

The authors declare no competing interests.

### Additional Information

Supplementary Information is available for this paper: supplementary data are contained in Extended Data Figures 1-7, supplementary methodologic information in Tables 1-3, and raw data for all figures generated in this work in Extended Data File 1. Correspondence and requests for materials should be directed to Rachel Perry. Reprints and permissions information is available at www.nature.com/reprints.

## Extended Data

**Extended Data Figure 1.**
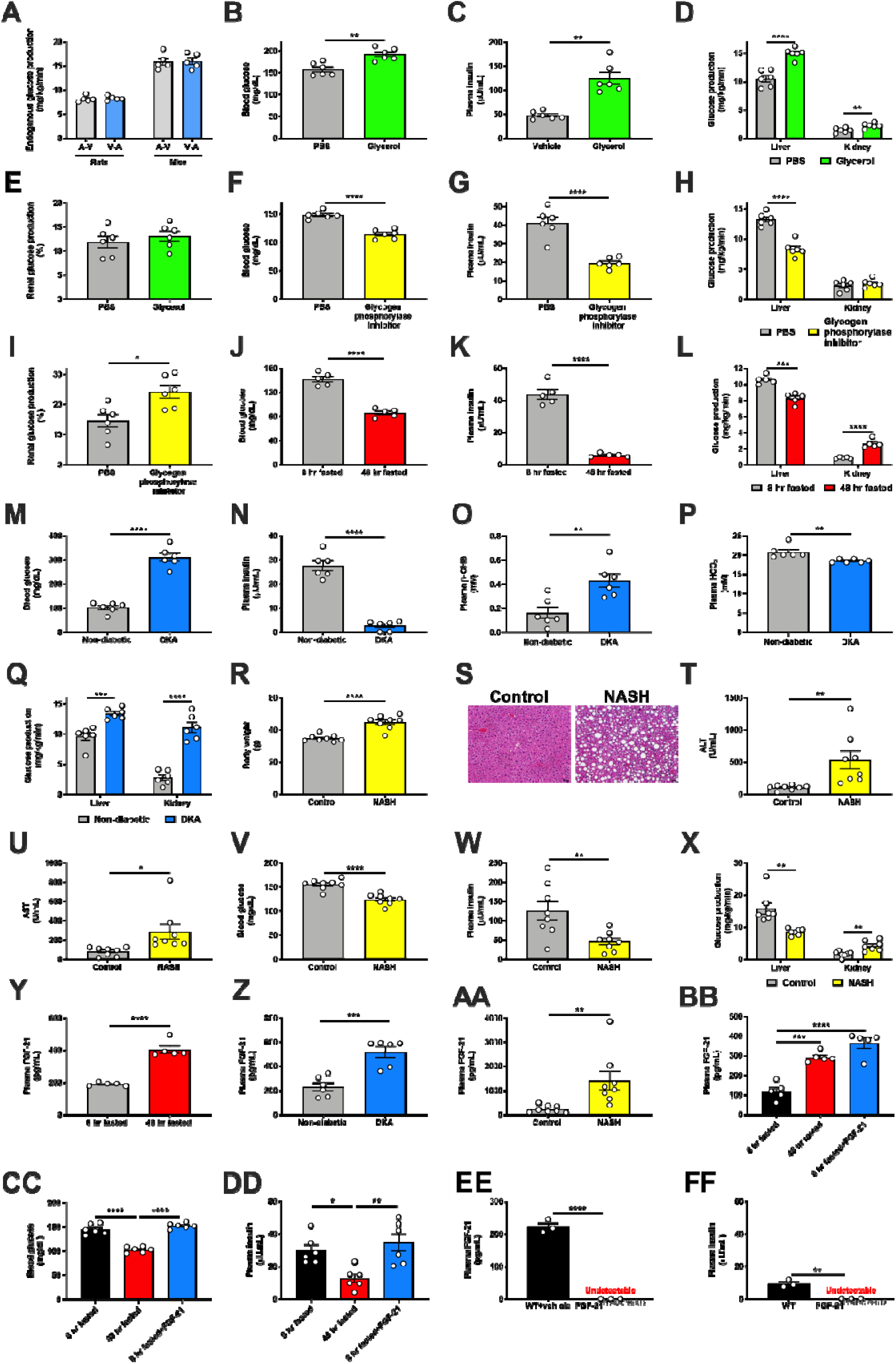
FGF-21 promotes renal gluconeogenesis under conditions of metabolic stress. (A) Measured endogenous glucose production is identical in rats and mice whether tracer is infused into the carotid artery and blood drawn from the jugular vein (A-V), or tracer is infused into the jugular vein and blood drawn from the carotid artery (V-A) (n=5 per group). (B)-(C) Blood glucose and plasma insulin concentrations in 6 hr fasted mice infused with glycerol, a gluconeogenic substrate (n=6 per group). (D)-(E) Hepatic and renal glucose production (n=6 per group). (F)-(G) Blood glucose and plasma insulin in 6 hr fasted mice treated with a glycogen phosphorylase antagonist to inhibit glycogenolysis (n=6 per group). (H)-(I) Hepatic and renal glucose production (n=6 per group). (J)-(K) Blood glucose and plasma insulin in recently fed (8 hr fasted) and starved (48 hr fasted) mice (n=5 per group). (L) Hepatic and renal glucose production (n=5 per group). (M)-(P) Blood glucose, plasma insulin, β-OHB, and bicarbonate in a mouse model of diabetic ketoacidosis (n=6 per group). (Q) Hepatic and renal glucose production (n=6 per group). (R)-(S) Body weight and liver hematoxylin & eosin staining in a mouse model of NASH (n=8 per group). Scale bar, 100 μm. (T)-(U) Liver transaminase concentrations (n=8 per group). (V)-(W) Blood glucose and plasma insulin (n=8 per group). (X) Hepatic and renal glucose production (n=7 per group). (Y)-(AA) Plasma FGF-21 concentrations in fed/fasted (n=5 per group), DKA (n=6 per group), and NASH models (n=8 per group). (BB)-(DD) Plasma FGF-21 (n=5 per group), blood glucose (n=6 per group), and plasma insulin concentrations (n=6 per group) in recently fed, fasted, and FGF-21 infused rats. \**P*<0.05, \*\**P*<0.01, \*\*\**P*<0.001, \*\*\*\**P*<0.0001 by ANOVA with Tukey’s multiple comparisons test. (EE)-(FF) Plasma FGF-21 and insulin concentrations in FGF-21^f/f;Alb-CreERT2^ mice and their WT littermates fasted for 48 hr (n=3 per group). In all panels, \**P*<0.05, \*\**P*<0.01, \*\*\**P*<0.001, \*\*\*\**P*<0.0001 by the 2-tailed unpaired Student’s t-test.

**Extended Data Figure 2.**
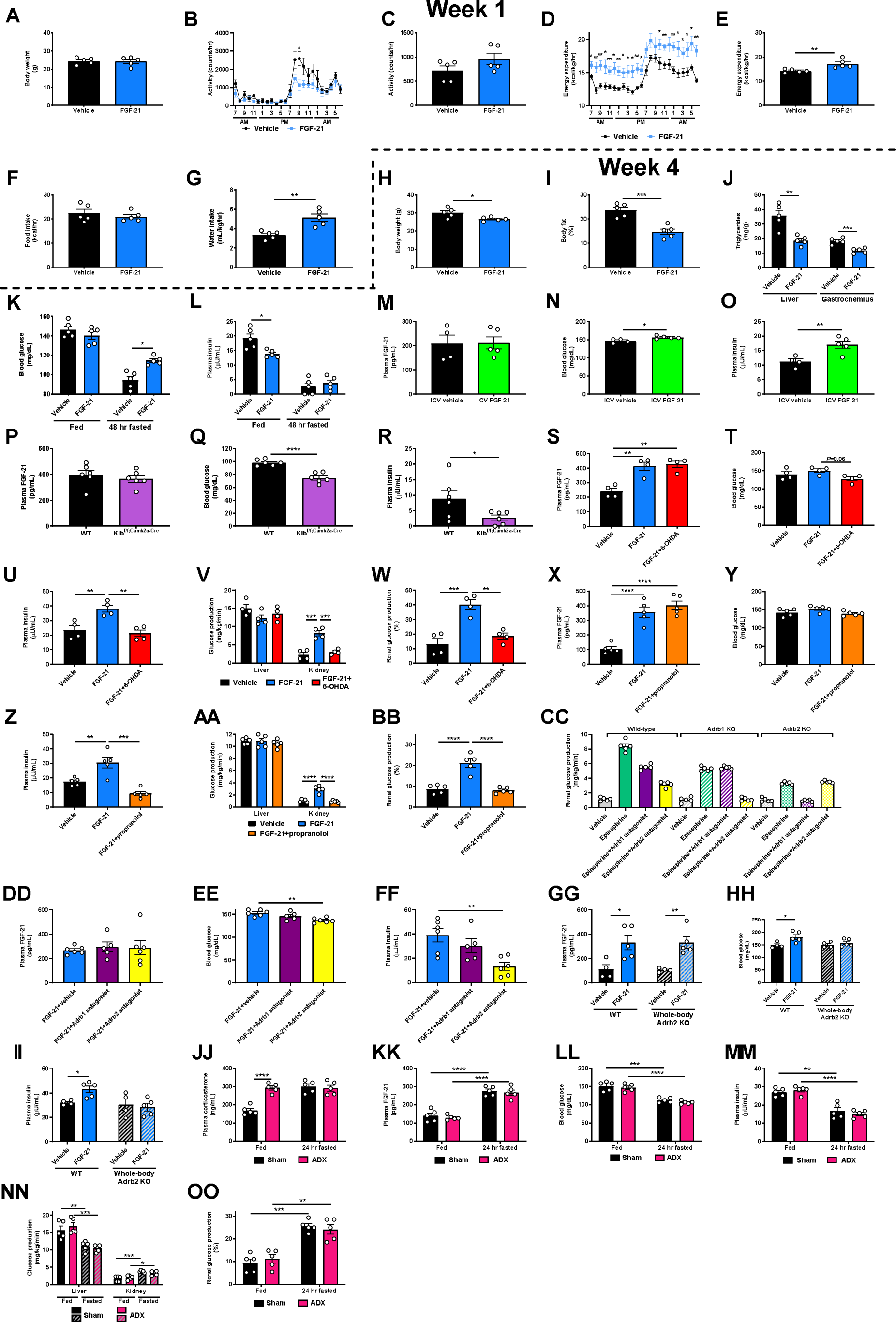
FGF-21 promotes renal gluconeogenesis via Adrb2-dependent neural hardwiring. Consistent with previous reports, we find that chronic FGF-21 infusion increases energy expenditure and improves metabolic health in diet-induced obese mice. (A)-(G) Body weight, activity, energy expenditure, food and water intake during the first week of FGF-21 or vehicle infusion. Throughout this figure, unless otherwise specified, groups were compared by the 2-tailed unpaired Student’s t-test. In panels (A)-(L), n=5 per group. (H)-(L) Body weight and fat, tissue triglyceride content, blood glucose, and plasma insulin concentrations in week 4 of FGF-21 infusion. (M) Jugular vein plasma FGF-21 concentrations in rats infused with FGF-21 into the third ventricle (ICV) (n=4 vehicle-treated and 5 FGF-21-treated rats in panels (M)-(O)). (N)-(O) Blood glucose and plasma insulin. (P)-(R) Plasma FGF-21, blood glucose, and plasma insulin concentrations in WT and Klb^f/f;Camk2a-Cre^ mice (n=6 per group). (S)-(U) Plasma FGF-21, blood glucose, and plasma insulin concentrations in FGF-21 infused mice, chemically sympathectomized with 6-OHDA (n=4 per group). In panels (S)-(FF), groups were compared by ANOVA with Tukey’s multiple comparisons test. (V)-(W) Hepatic and renal glucose production (n=4 per group). (X)-(Z) Plasma FGF-21, blood glucose, and plasma insulin concentrations in mice infused with FGF-21 and treated with the nonselective Adrb antagonist propranolol (n=5 per group). (AA)-(BB) Hepatic and renal glucose production (n=5 per group). (CC) Validation of Adrb1 and Adrb2 antagonists: epinephrine-stimulated renal glucose production (n=5 per group). For clarity of presentation, statistical comparisons were not performed. (DD) Plasma FGF-21 concentrations in FGF-21 infused mice treated with antagonists of Adrb1 (betaxolol) or Adrb2 (butoxamine) (in panels (DD)-(FF), n=6 [vehicle and Adrb2 antagonist-treated] or 5 per group [Adrb1 antagonist-treated]). (EE) Blood glucose. (FF) Plasma insulin concentrations. (GG) Plasma FGF-21 concentrations in vehicle- and FGF-21-infused WT and whole-body Adrb2 KO littermates (in panels (GG)-(II), n=4 (vehicle-treated) or 5 (FGF-21-treated) per group. (HH)-(II) Blood glucose and plasma insulin concentrations. (JJ) Plasma corticosterone in sham-operated and adrenalectomized mice. ADX mice were implanted with a subcutaneous pump to deliver corticosterone to match concentrations in 24 hr fasted mice, in order to avoid corticosterone as a potential phenotypic confounder. (KK) Plasma FGF-21. (LL)-(MM) Hepatic and renal glucose production. (NN)-(OO) Blood glucose and plasma insulin concentrations. In panels (JJ)-(OO), n=5 per group, and fed vs. 24 hr fasted and sham vs. ADX mice were compared by the 2-tailed unpaired Student’s t-test. In all panels, \**P*<0.05, \*\**P*<0.01, \*\*\**P*<0.001, \*\*\*\**P*<0.0001.

**Extended Data Figure 3.**
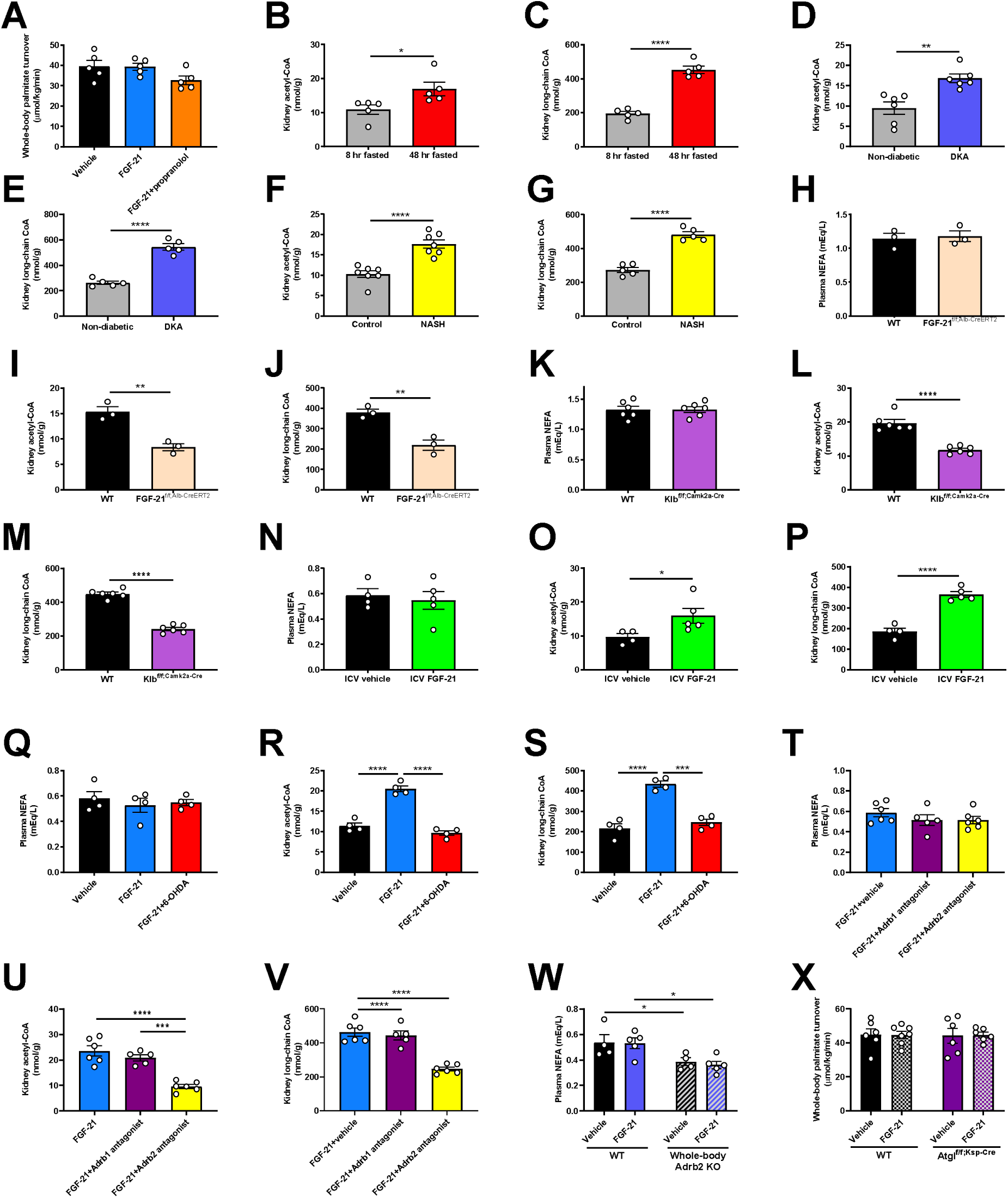
FGF-21-induced increases in renal lipolysis promote increased renal gluconeogenesis in metabolic stress. (A) Whole-body lipolysis (palmitate turnover) in mice infused with FGF-21±pretreatment with propranolol. In panels (A), (Q)-(V), groups were compared by ANOVA with Tukey’s multiple comparisons test. In panels (A)-(C), n=5 per group. (B)-(C) Kidney acetyl- and long-chain acyl-CoA concentrations in fed/fasted mice. (D)-(E) Kidney acetyl- and long-chain acyl-CoA concentrations in mice in DKA (n=6 per group). (F)-(G) Kidney acetyl- and long-chain acyl-CoA concentrations in a mouse model of NASH (n=8 per group). (H) Plasma non-esterified fatty acid concentrations in 24 hr fasted FGF-21^f/f;Alb-CreERT2^ mice (i.e. liver-specific FGF-21 knockout) (n=3 per group). (I)-(J) Kidney acetyl- and long-chain acyl-CoA concentrations. (K) Plasma NEFA in Klb^f/f;Camk2a-Cre^ mice (i.e. brain-specific Klb knockout). In panels (K)-(M), n=5 per group. (L)-(M) Kidney acetyl- and long-chain acyl-CoA concentrations (n=5 per group). (N) Plasma NEFA in ICV FGF-21 infused rats. In panels (N)-(P), n=4 vehicle-treated and 5 FGF-21-treated rats per group. (O)-(P) Kidney acetyl- and long-chain acyl-CoA concentrations. (Q) Plasma NEFA in vehicle and FGF-21-infused mice, some pre-treated with chemical sympathectomy via 6-OHDA (in panels (Q)-(S), n=4 per group). (R)-(S) Kidney acetyl- and long-chain acyl-CoA concentrations. (T) Plasma NEFA in FGF-21-infused mice, some treated with antagonists of Adrb1 or Adrb2 (in panels (T)-(V), n=6 vehicle- or Adrb2 antagonist-treated mice, or 5 Adrb1 antagonist-treated mice). (U)-(V) Kidney acetyl- and long-chain acyl-CoA concentrations. (W) Plasma NEFA in WT and whole-body Adrb2 KO mice (n=4 vehicle-treated or 5 FGF-21-treated mice per group). (X) Whole-body palmitate turnover in Atgl^f/f;Ksp-Cre^ mice infused with FGF-21 or vehicle, and their WT littermates (n=6 per group, with the exception of WT+FGF-21-treated mice, in which n=7 per group). Unless otherwise specified, groups were compared by the 2-tailed unpaired Student’s t-test. In all panels, \**P*<0.05, \*\**P*<0.01, \*\*\**P*<0.001, \*\*\*\**P*<0.0001.

**Extended Data Figure 4.**
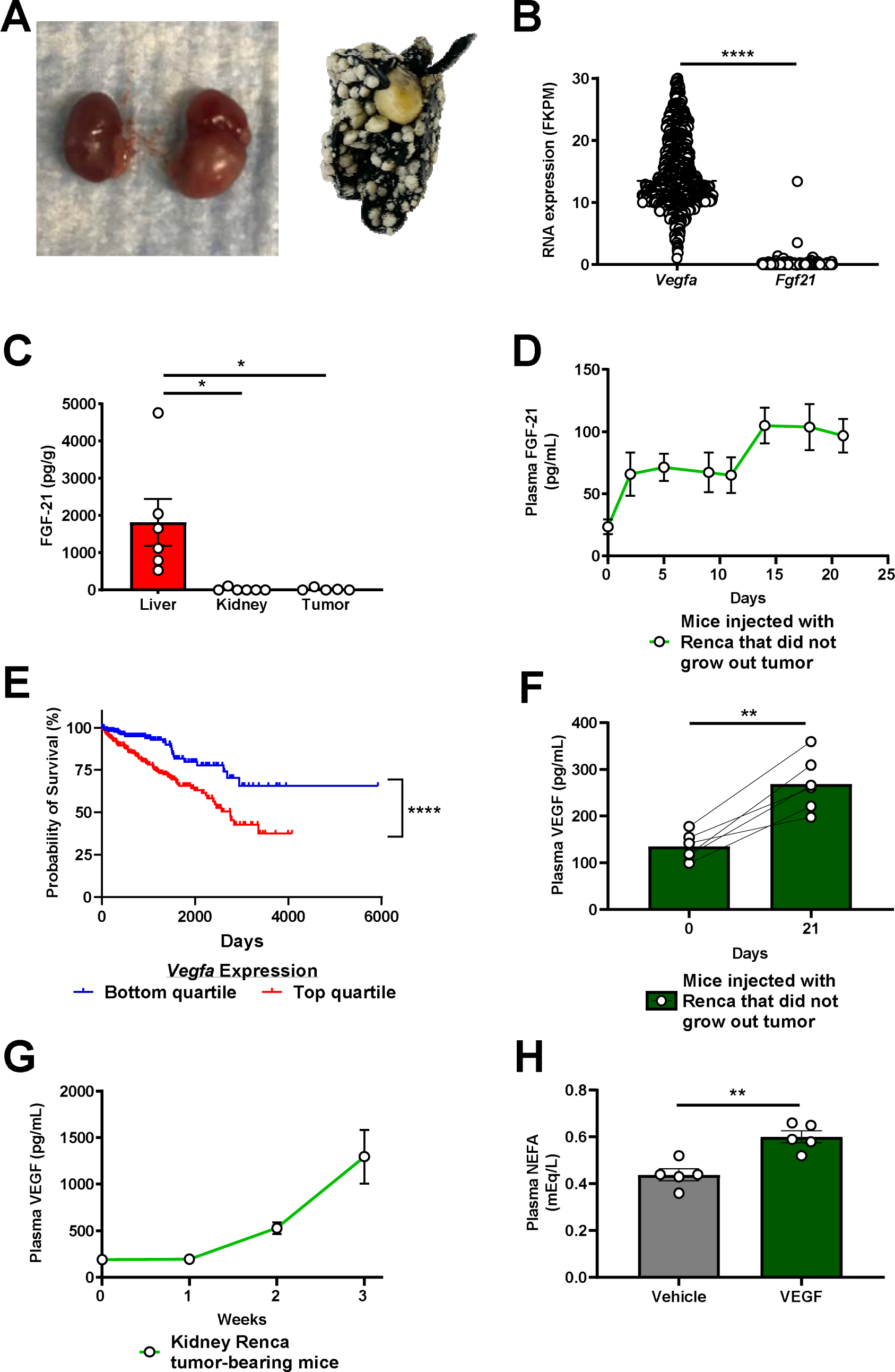
FGF-21 is increased in renal cell carcinoma due to increased circulating VEGF. (A) Photos of kidneys from the same mouse, injected with PBS (left) or with Renca cells (right), and a lung from a Renca tumor-bearing mouse, stained with India ink. Metastases appear white. (B) Tumor *Vegfa* and *Fgf21* mRNA expression in human renal cell carcinoma (n=877 for both proteins). Data from the Human Protein Atlas^43^. FKPM, fragments per kilobase of transcript per million mapped reads. (C) Tissue FGF-21 protein, measured by ELISA (n=6 liver, 6 kidney, and 5 tumor). (D) Plasma FGF-21 concentrations in mice injected with Renca cells into the renal cortex that did not ultimately grow out a palpable tumor (n=5 on day 0, and 6 on all subsequent days). (E) Survival of RCC patients whose tumors were in the upper and lower quartile (n=219 per quartile) for *Vegfa* expression. (F) Plasma VEGF concentrations in mice injected with Renca cells into the renal cortex that did not ultimately grow out a visualized tumor (n=6). (G) Plasma VEGF concentrations in mice with Renca RCC tumors (n=5 per timepoint until week 3, at which n=4). (H) Plasma NEFA concentrations in mice injected with recombinant VEGF (n=5 per group). In all panels, \**P*<0.05, \*\**P*<0.01, \*\*\**P*<0.001, \*\*\*\**P*<0.0001.

**Extended Data Figure 5.**
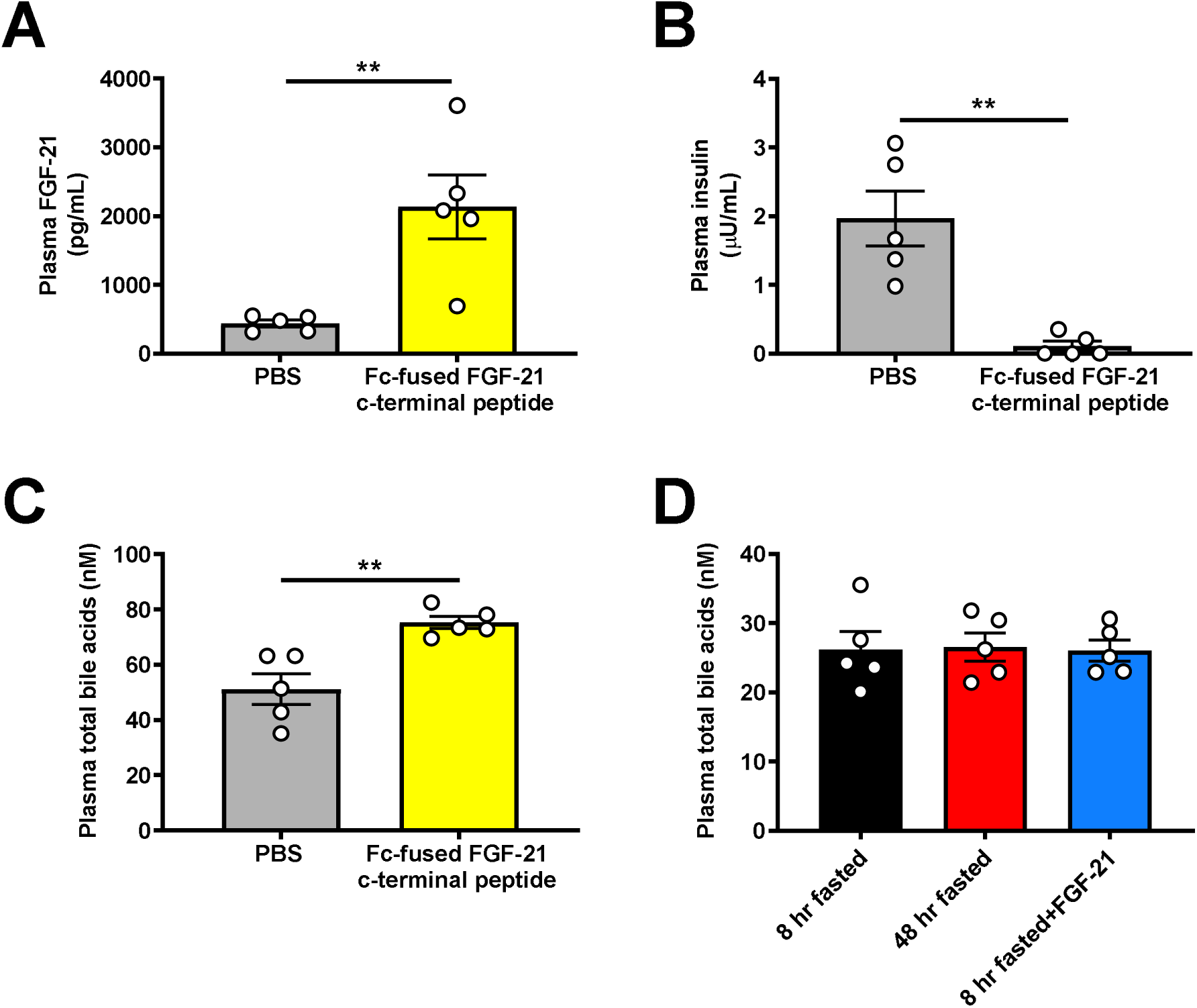
FGF-21 promotes renal glucose production in mice with renal cell carcinoma. (A) Plasma FGF-21, (B) Plasma insulin, and (C) Plasma total bile acids in mice treated with an Fc-fused FGF-21 c-terminal peptide. (D) Plasma total bile acids in rats fasted for 8 or 48 hours, and 8 hr fasted rats infused with FGF-21. In all panels, n=5 per group. \**\*P<*0.01.

**Extended Data Figure 6.**
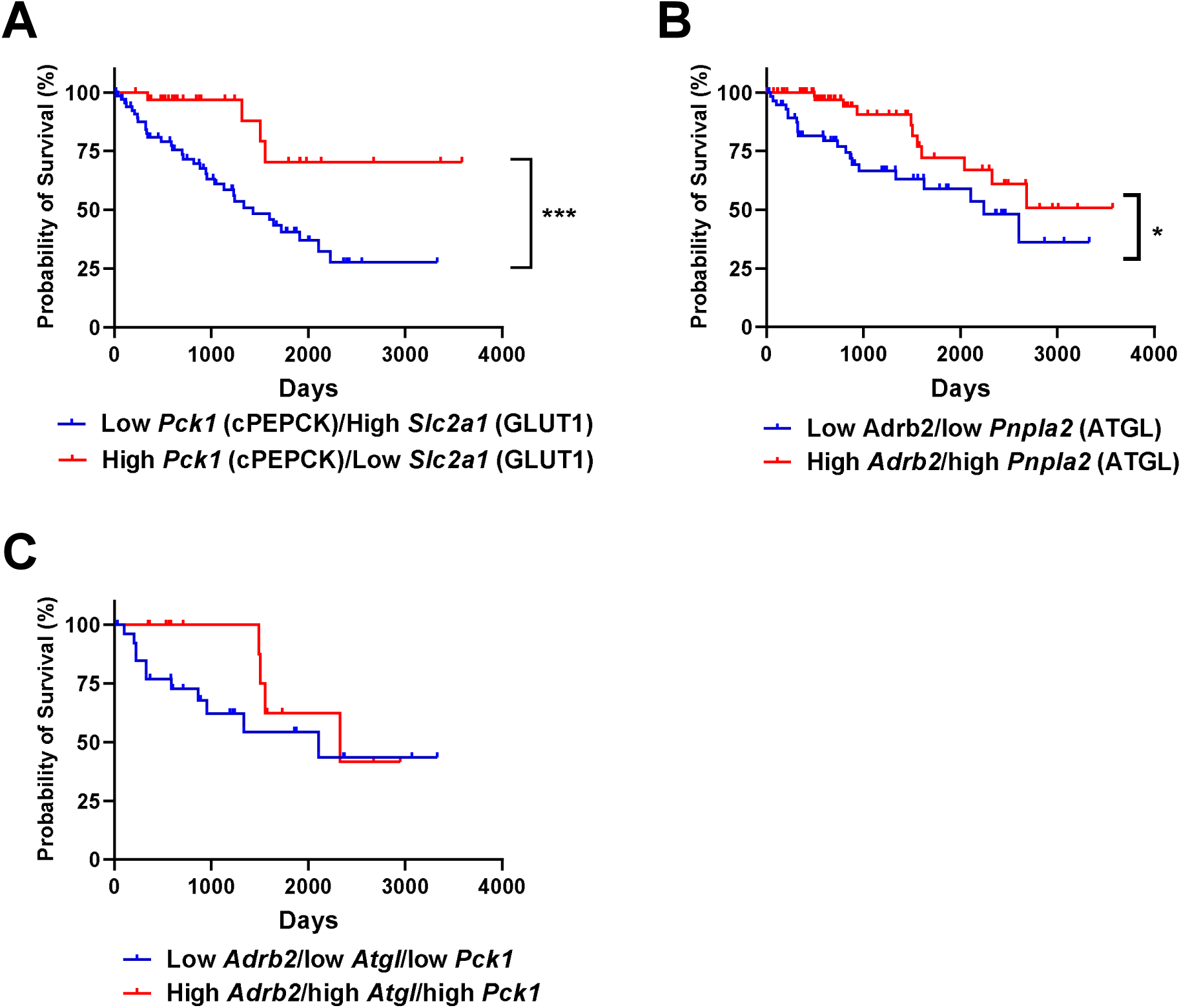
Renal gluconeogenesis is a targetable, pathogenic factor in murine models of RCC. (A) Patients with low *Slc2a1* and high cytosolic *Pck1* expression (i.e. low glucose uptake through GLUT1 and high glucose production facilitated by PC expression at the transcriptional level) have poorer survival as compared to patients with high *Slc2a1* and low *Pck1* expression in tumor^43^ (n=67 low *Slc2a1* and high *Pck1*, vs. 32 high *Slc2a1* and low *Pck1*). Unfortunately, expression data from surrounding parenchyma are not available. In panels (A) and (B), “low” and “high” were defined as falling in both the upper and lower, or lower and upper quartile of expression of the genes of interest. (B) Patients with low *Atgl (Pnpla2)* and *Adrb2* expression in RCC tumors (n=60) have worse survival than patients with high *Atgl (Pnpla2)* and *Adrb2* expression in RCC tumors (n=82). (C) Survival of RCC patients whose tumors express low *Adrb2*, low *Atgl*, and low *Pck 1* (n=27) vs. those whose tumors express high *Adrb2*, high *Atgl*, and high *Pck 1* (n=13).

**Extended Data Figure 7.**
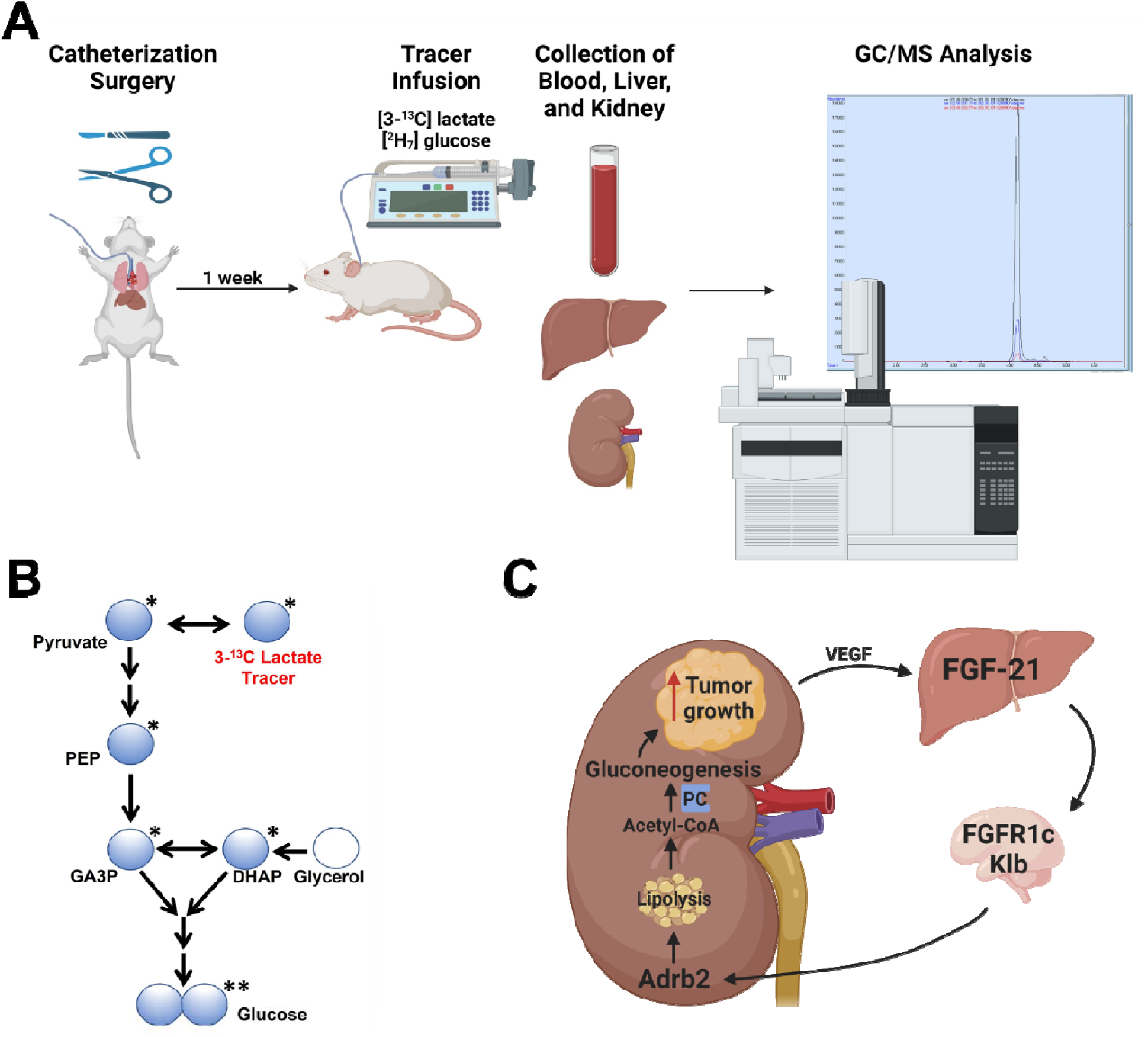
REGAL workflow and mechanistic summary. (A) REGAL workflow. Figure created with BioRender.com and modified to add a GC/MS spectrum generated by the authors. (B) Mass Isotopomer Distribution Analysis strategy. (C) Proposed mechanism by which FGF-21 promotes renal glucose production and, in turn, renal cell carcinoma. Figure created with BioRender.com.

**Extended Data Table 1.** **Clinical characteristics** of patients whose RCC tumor and kidney parenchyma samples were analyzed. Samples from 14 patients were studied; however, not all clinical data were available for all patients.

| Parameter | Fraction or Average±SEM |
| --- | --- |
| Clear cell RCC/other RCC | 7/7 |
| Stage | 1.5±0.3 |
| Necrosis (+/-) | 6/6 |
| Age (years) | 64±4 |
| Sex (F/M) | 6/7 |

